# A mind-body interface alternates with effector-specific regions in motor cortex

**DOI:** 10.1101/2022.10.26.513940

**Authors:** Evan M. Gordon, Roselyne J. Chauvin, Andrew N. Van, Aishwarya Rajesh, Ashley Nielsen, Dillan J. Newbold, Charles J. Lynch, Nicole A. Seider, Samuel R. Krimmel, Kristen M. Scheidter, Julia Monk, Ryland L. Miller, Athanasia Metoki, David F. Montez, Annie Zheng, Immanuel Elbau, Thomas Madison, Tomoyuki Nishino, Michael J. Myers, Sydney Kaplan, Carolina Badke D’Andrea, Damion V. Demeter, Matthew Feigelis, Deanna M. Barch, Christopher D. Smyser, Cynthia E. Rogers, Jan Zimmermann, Kelly N. Botteron, John R. Pruett, Jon T. Willie, Peter Brunner, Joshua S. Shimony, Benjamin P. Kay, Scott Marek, Scott A. Norris, Caterina Gratton, Chad M. Sylvester, Jonathan D. Power, Conor Liston, Deanna J. Greene, Jarod L. Roland, Steven E. Petersen, Marcus E. Raichle, Timothy O. Laumann, Damien A. Fair, Nico U.F. Dosenbach

**Affiliations:** Mallinckrodt Institute of Radiology, Washington University School of Medicine, St. Louis, MO, 63110, USA; Department of Neurology, Washington University School of Medicine, St. Louis, MO, 63110, USA; Department of Biomedical Engineering, Washington University in St. Louis, St. Louis, MO, 63130, USA; Department of Neurology, New York University Langone Medical Center, New York, NY 10016, USA; Department of Psychiatry, Weill Cornell Medicine, New York, NY, 10021, USA; Department of Psychiatry, Washington University School of Medicine, St. Louis, MO, 63110, USA; Department of Pediatrics, University of Minnesota, Minneapolis, MN, 55454, USA; Department of Cognitive Science, University of California San Diego, La Jolla, CA 92093; Department of Psychological and Brain Sciences, Washington University in St. Louis, St. Louis, MO, 63130, USA; Department of Pediatrics, Washington University School of Medicine, St. Louis, MO, 63110, USA; Department of Neuroscience, University of Minnesota, Minneapolis, MN, 55454, USA; Department of Neurosurgery, Washington University School of Medicine, St. Louis, MO, 63110, USA; Department of Psychology, Florida State University, Tallahassee, FL, 32306, USA; Department of Neuroscience, Washington University School of Medicine, St. Louis, MO, 63110, USA; Masonic Institute for the Developing Brain, University of Minnesota, Minneapolis, MN, 55454, USA; Program in Occupational Therapy, Washington University in St. Louis, St. Louis, MO, 63130, USA

## Abstract

Primary motor cortex (M1) has been thought to form a continuous somatotopic homunculus extending down precentral gyrus from foot to face representations^1,2^. The motor homunculus has remained a textbook pillar of functional neuroanatomy, despite evidence for concentric functional zones^3^ and maps of complex actions^4^. Using our highest precision functional magnetic resonance imaging (fMRI) data and methods, we discovered that the classic homunculus is interrupted by regions with sharpy distinct connectivity, structure, and function, alternating with effector-specific (foot, hand, mouth) areas. These inter-effector regions exhibit decreased cortical thickness and strong functional connectivity to each other, and to prefrontal, insular, and subcortical regions of the Cingulo-opercular network (CON), critical for executive action^5^ and physiological control^6^, arousal^7^, and processing of errors^8^ and pain^9^. This interdigitation of action control-linked and motor effector regions was independently verified in the three largest fMRI datasets. Macaque and pediatric (newborn, infant, child) precision fMRI revealed potential cross-species analogues and developmental precursors of the inter-effector system. An extensive battery of motor and action fMRI tasks documented concentric somatotopies for each effector, separated by the CON-linked inter-effector regions. The inter-effector regions lacked movement specificity and co-activated during action planning (coordination of hands and feet), and axial body movement (e.g., abdomen, eyebrows). These results, together with prior work demonstrating stimulation-evoked complex actions^4^ and connectivity to internal organs (e.g., adrenal medulla)^10^, suggest that M1 is punctuated by an integrative system for implementing whole-body action plans. Thus, two parallel systems intertwine in motor cortex to form an integrate-isolate pattern: effector-specific regions (foot, hand, mouth) for isolating fine motor control, and a mind-body interface (MBI) for the integrative whole-organism coordination of goals, physiology, and body movement.

## MAIN

Beginning in the 1930s, Penfield and colleagues mapped human M1 with direct cortical stimulation, eliciting movements from about half of sites, mostly of the foot, hand, or mouth^1^. Although representations for specific body parts overlapped substantially^11^, these maps gave rise to the textbook view of M1 organization as a continuous homunculus, from head to toe.

In non-human primates, organizational features inconsistent with the motor homunculus have been described. Structural connectivity studies divided M1 into anterior, gross-motor, “old” M1 (few direct projections to spinal motoneurons), and posterior, fine-motor, “new” M1 (many direct motoneuronal projections)^12,13^. Non-human primate stimulation studies showed the body to be represented in anterior M1^14^, and the motor effectors (tail, foot, hand, mouth) in posterior M1. Such studies also suggested that the limbs are represented in concentric functional zones progressing from the digits at the center, to the shoulders on the periphery ^3,15–17^. Moreover, stimulations could elicit increasingly complex and multi-effector actions when moving from posterior to anterior M1^4,18,19^.

During natural behavior, voluntary movements are part of goal-directed actions, initiated and controlled by executive regions in the CON^5,20^. Neural activity preceding voluntary movements can first be detected in the dorsal anterior cingulate cortex (dACC) or rostral cingulate zone^21^, then in the pre-supplementary motor area (pre-SMA) and SMA^22^, followed by M1. These regions all project to the spinal cord^23^, with M1 as the main transmitter of motor commands down the corticospinal tract^24^. Efferent motor copies are received by primary somatosensory cortex (S1)^25^, cerebellum, and striatum^26^ for online correction and learning. Tracer injections in non-human primates demonstrated direct projections from anterior M1/CON to internal organs (e.g. adrenal medulla) for preparatory sympathetic arousal in anticipation of action^10^. Post-movement error and pain signals are relayed primarily to insular and cingulate regions of the CON, which update future action plans^8,9^.

Resting state functional connectivity (RSFC) fMRI noninvasively maps the brain’s functional networks^27,28^. Precision functional mapping (PFM) studies rely on large amounts of multi-modal data (e.g., RSFC, tasks) to map individual-specific brain organization in greatest possible detail^29–31^. Early PFM studies identified separate foot, hand, and mouth M1 regions^23^ with their respective cerebellar and striatal targets^26,27^. These foot/hand/mouth motor circuits were characterized by strong within-circuit connectivity and effector specificity in task fMRI^23^. However, these circuits were relatively isolated and did not include functional connections with control networks such as CON that could support the integration of movement with global behavioral goals. A recent study showed that prolonged dominant arm immobilization strengthened functional connectivity between disused M1 and the CON^28,29^, suggesting that the CON’s role may extend beyond abstract action control and into movement coordination.

Here, we used the latest iteration of PFM with higher resolution (2.4 mm) and greater amounts of fMRI (RSFC: 172 – 1,813 min/participant; task: 353 min/participant), and diffusion data, to map M1 and its connections with highest detail. Results were verified in group-averaged data from the three largest fMRI studies (Human Connectome Project [HCP], Adolescent Brain Cognitive Development [ABCD] Study, UK Biobank [UKB]; total n ∼ 50,000). Furthermore, we placed our findings in cross-species (macaque vs human), developmental (neonate, infant, child, and adult), and clinical (perinatal stroke) contexts using PFM data.

### Two distinct functional systems alternate in motor cortex

Advanced PFM revealed connectivity that differed strikingly from the canonical homuncular organization of M1. Two contrasting patterns of functional connectivity alternated in primary motor cortex (Fig. 1a; Supplementary Movie 1). The expected pattern, as previously described for M1 foot, hand, and mouth representations^29,34^, was comprised of three regions (per hemisphere) for which cortical connectivity was restricted to homotopic contralateral M1, SMA and adjacent S1 (Fig. 1a, seeds 1, 3, 5). This set of RSFC-defined regions corresponded with task-evoked activity during foot, hand, and tongue movements (Fig. 1b; see Fig. S1 for other participants).

**Fig 1.**
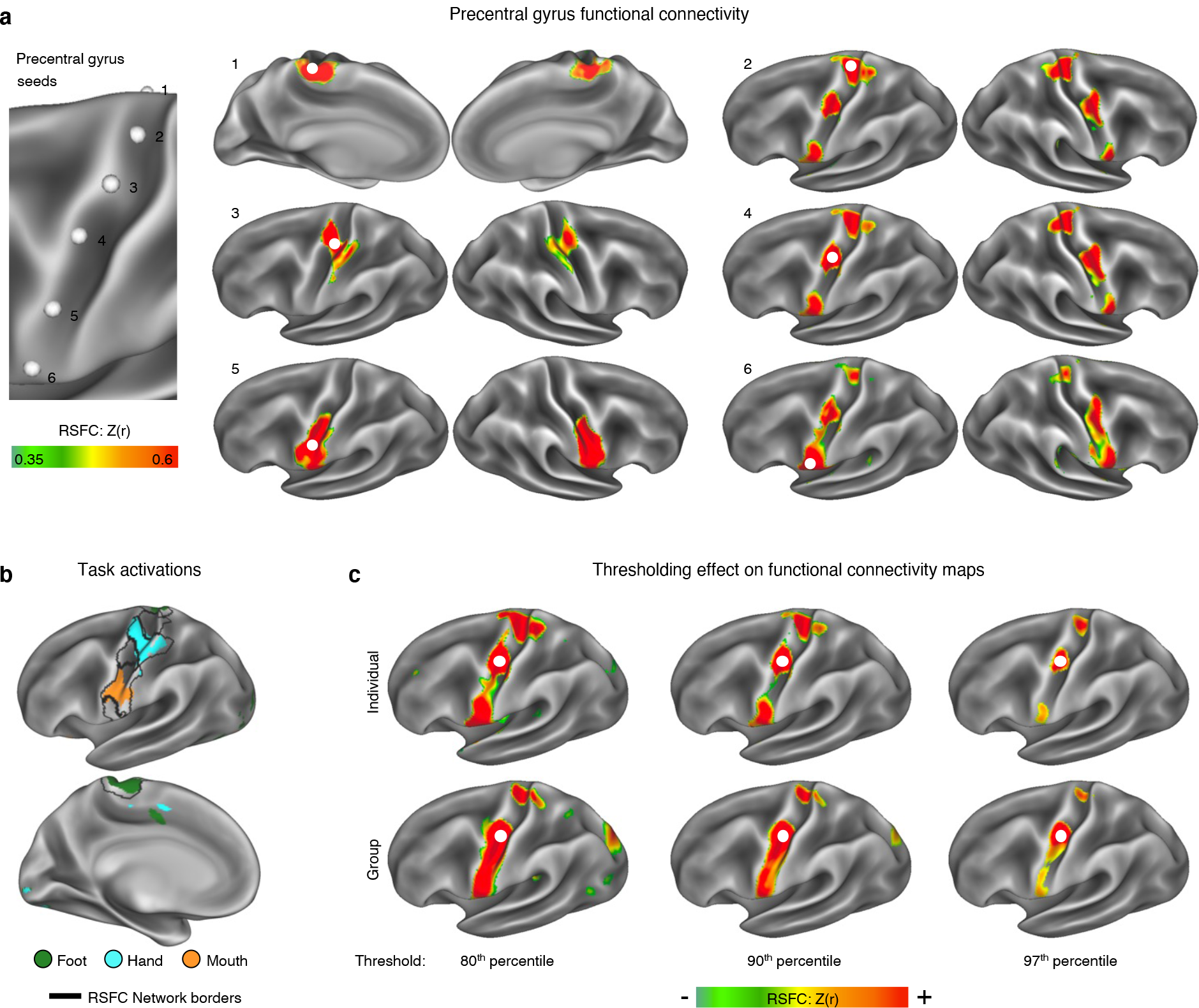
Precision functional mapping of primary motor cortex. **a**, Resting state functional connectivity (RSFC) seeded from a continuous line of cortical locations in the left precentral gyrus in a single exemplar participant (P1; 356 min resting-state fMRI). The six exemplar seeds shown represent all distinct connectivity patterns observed (see Supplementary Movie 1 for complete mapping). Functional connectivity seeded from these locations illustrated classical primary motor cortex connectivity of regions representing the foot (1), hand (3), and mouth (5), as well as an interdigitated set of strongly interconnected regions (2, 4, 6). See Fig. S1a and Supplementary Movie 2 for all highly-sampled participants, Fig. S1b for within-participant replications, and Fig. S1c for group-averaged data. **b**, Discrete functional networks were demarcated using a whole-brain, data-driven, hierarchical approach (see Methods) applied to the resting-state fMRI data, which defined the spatial extent of the networks observed in Fig. 1 (black outlines). Regions defined by RSFC were functionally labeled using a classic block-design fMRI motor task involving separate movement of the foot, hand, and tongue (following^34^; see^32^ for details). The map illustrates the top 1% of vertices activated by movement of the foot (green), hand (cyan), and mouth (orange) in the exemplar participant (P1; see Fig. S1d for other participants). **c**, The inter-effector connectivity pattern became more distinct from surrounding effector-specific motor regions as connectivity thresholding increased from the 80^th^ to the 97^th^ percentile. RSFC thresholds required to detect the inter-effector pattern were lower in individual-specific data (top) than in group-averaged data (ABCD Study, bottom).

Interleaved between the known foot/hand/mouth M1 regions lay three areas that were strongly functionally connected to each other, both contralaterally and ipsilaterally, forming a previously unrecognized interdigitated chain down the precentral gyrus (Fig. 1a, seeds 2, 4, 6). The motif of three M1 inter-effector regions was observed in every highly-sampled adult (Fig. S1a; Table S1) and replicated within-individual in separate data from the same participants (Fig. S1b). Importantly, the inter-effector pattern was also evident in all large-N group-average datasets (UKB, ABCD, HCP, WU120; Fig. S1c). The M1 inter-effector functional connectivity motif was most apparent in individual-specific maps, but once recognized, was also clearly identifiable in group-averaged data when visualized using stringent connectivity thresholds (Fig. 1c).

The inter-effector regions were evident relatively early in development. While PFM data from a newborn failed to reveal the inter-effector motif, it was detectable in an 11-month-old infant, and was almost adult-like in a 9-year-old child (Fig. S2a-e). Inter-effector regions could even be identified in an individual with preserved motor function despite suffering severe bilateral perinatal strokes that destroyed large portions of M1 (Fig. S2f; see ^35^ for clinical details).

### Inter-effector regions are functionally connected to Cingulo-opercular control network

In addition to being interconnected, the three inter-effector regions were functionally connected to the dACC and pre-SMA, thought to be important for goal-oriented cognitive control. In striatum, inter-effector regions were most strongly connected to dorsolateral putamen. In thalamus, connectivity peaked in the centromedian (CM) nucleus (Fig. 2a; see Fig. S3, Table S1 for other participants), with additional strong connectivity observed in ventral intermediate (VIM), ventral posteriomedial (VPM), and ventral posterior inferior (VPI) nuclei. Inter-effector regions were strongly connected to cerebellar areas surrounding but distinct from effector-specific cerebellar regions (Fig. S1e).

**Fig 2.**
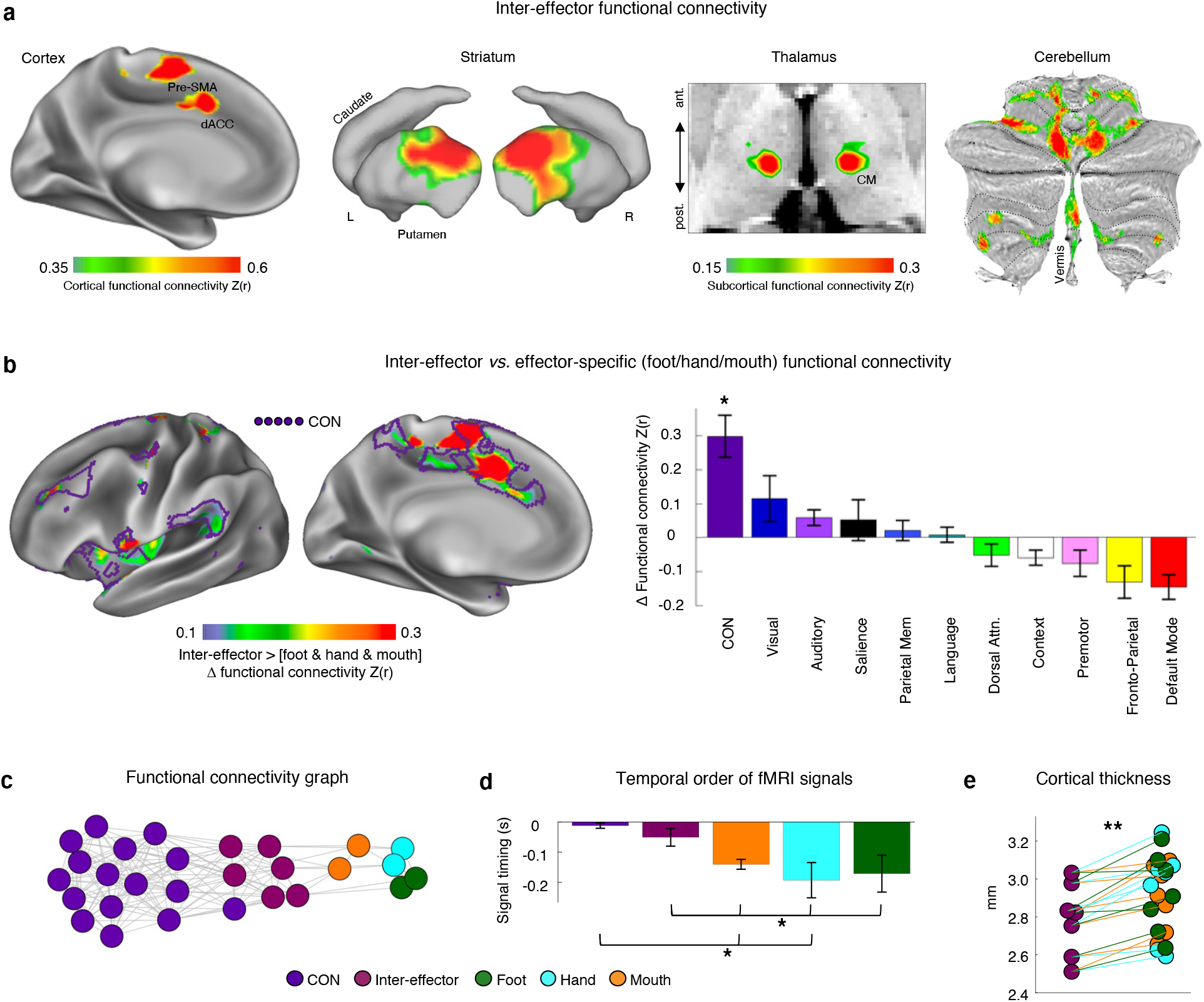
Functional connectivity and cortical thickness of the motor cortex inter-effector motif. **a**, Brain regions with the strongest functional connectivity to the left middle inter-effector region (exemplar seed) in cortex, striatum, thalamus (horizontal slice; centromedian (CM) nucleus shown), and cerebellum (flat map) in the exemplar participant (P1); Fig. S3 for other participants. **b**, Brain regions more strongly functionally connected to inter-effectors than to any foot/hand/mouth regions (P1; Fig. S4a for other participants). Purple outlines show the Cingulo-opercular Network (CON; individual-specific). Central sulcus is masked as it exhibits large differences by definition. **c**, Connectivity was calculated between every network and both inter-effector and effector-specific M1 regions. The plot shows the smallest difference between inter-effector connectivity and any effector-specific connectivity (standard error bars). This difference was larger for CON than for any other network (*: *P* < 0.05). **c**, Inter-network relationships visualized in network space using a spring-embedding plot, in which connected regions are pulled together while disconnected regions are pushed apart. Connecting lines indicate a functional connection (r > 0.2) (P1; Fig. S4b for all participants). **d**, Inter-effector and effector-specific regions were tested for systematic differences in the temporal ordering of their infra-slow fMRI signals (<0.1 Hz)^36^. Plot shows across-participant average signal ordering in CON, inter-effector, and effector-specific regions (standard error bars; *: *P* < 0.05). Prior electrophysiology work suggests that later infra-slow activity (here, CON) corresponds to earlier delta-band (0.5-4Hz) activity^37^. **e**, In each participant (individual dots), inter-effector regions exhibited lower cortical thickness than all effector-specific regions (**: *P* < 0.01).

In all highly-sampled individuals (n = 7), the inter-effector regions had stronger connections to CON than did any of the foot/hand/mouth regions (Fig. 2b; Fig. S4a for all participants; across participants: all paired t > 4.75; all *P* < 0.01; Fig. S5a). The inter-effector *vs*. foot/hand/mouth difference was larger for CON than for any other network (all paired t > 2.8; all *P* < 0.03; Fig. 2b). In network space, inter-effector regions were positioned between CON and the foot/hand/mouth regions (Fig. 2c; Fig. S4b for all participants). Inter-effector regions were also more strongly connected to: middle insula, known to process pain^9^ and interoceptive^38^ signals (Fig. S5b; all paired t > 2.7; all *P* < 0.03); lateral cerebellar lobule V and vermis Crus II, lobule VIIb, and lobule VIIIa (all paired t > 3.7, all *P* < 0.01; Fig. S5c); dorsolateral putamen, critical for motor function (all paired t > 3.7; all *P* < 0.01, Fig. S5d); and sensory-motor regions of thalamus (VIM; CM; VPM; all paired t > 3.0, all *P* < 0.02).

Comparing the relative timing of resting-state fMRI signals (lag structure^36^) showed that infra-slow (<0.1 Hz) fMRI signals in both the CON and the inter-effector network lagged behind those in effector-specific regions (Fig. 2d; CON vs foot: paired t = 2.38, *P* = 0.055; vs hand and mouth: all t > 2.84, all *P* < 0.03; inter-effector vs foot/hand/mouth: all t > 2.5, all *P* < 0.05). Inter-regional lags in infra-slow (<0.1 Hz) signals are associated with propagation of higher-frequency delta activity (0.5-4 Hz) in the opposite direction^37^, suggesting that high-frequency signals may occur earlier in CON than M1—consistent with electrical recordings during voluntary movement^39^—but that such signals reach the inter-effectors earlier than the foot, hand, mouth regions.

As expected, the M1 foot/hand/mouth regions were strongly functionally connected with adjacent S1 (Fig. 1a, Fig. S6a), consistent with known functional connections between M1 and S1^28^. By contrast, inter-effector regions exhibited lower connectivity with adjacent S1 (Fig. S5h; all paired t > 3.2, all *P* < 0.02). More specifically, inter-effector functional connectivity extended into the fundus of the central sulcus (Fig. S6b; Brodmann Area [BA] 3a), which represents proprioception^40^, but not to the postcentral gyrus (BAs 3b/1/2) representing cutaneous tactile stimuli.

Convergent with these functional differences, metrics of brain structure systematically differed between inter-effector and effector-specific regions. Inter-effector regions exhibited lower cortical thickness (all paired t > 3.6; all *P* ≤ 0.01; Fig. 2e), more similar to prefrontal cortex^41^, but higher fractional anisotropy (2 mm beneath cortex; all paired t > 5.3; all *P* < 0.05; Fig. S5j). Inter-effector regions also had higher intracortical myelin content than foot regions (paired t = 6.8, *P* < 0.001) but lower than hand regions (paired t = 4.8, *P* = 0.003; Fig. S5k).

### Concentric effector somatotopies are separated by inter-effector body/action regions

To better understand the functions of the inter-effector motif, we collected fMRI data during blocked performance of twenty-five different movements in two highly-sampled individuals (64 runs; 244 min/participant) and during a novel event-related task with separate planning and execution phases for coordinated hand and foot movements (12 runs; 132 min/participant). According to the homuncular model of M1, activation when moving a given body part should exhibit a single peak within the precentral gyrus. If M1 is instead organized into concentric functional zones, all movements except those at the centers (i.e., toes, fingers, tongue) should exhibit two peaks (above and below). Within each of the three effector-specific regions, the topography of preferred movements—the movement eliciting greatest activation in each vertex (Fig. 3a)—was more consistent with a concentric organization (distal-proximal; e.g.: toes in the center, with surrounding concentric zones of ankle-knee-hip)^3,42,43^ than with the canonical, linear toes-to-face homuncular model^1^.

**Fig 3.**
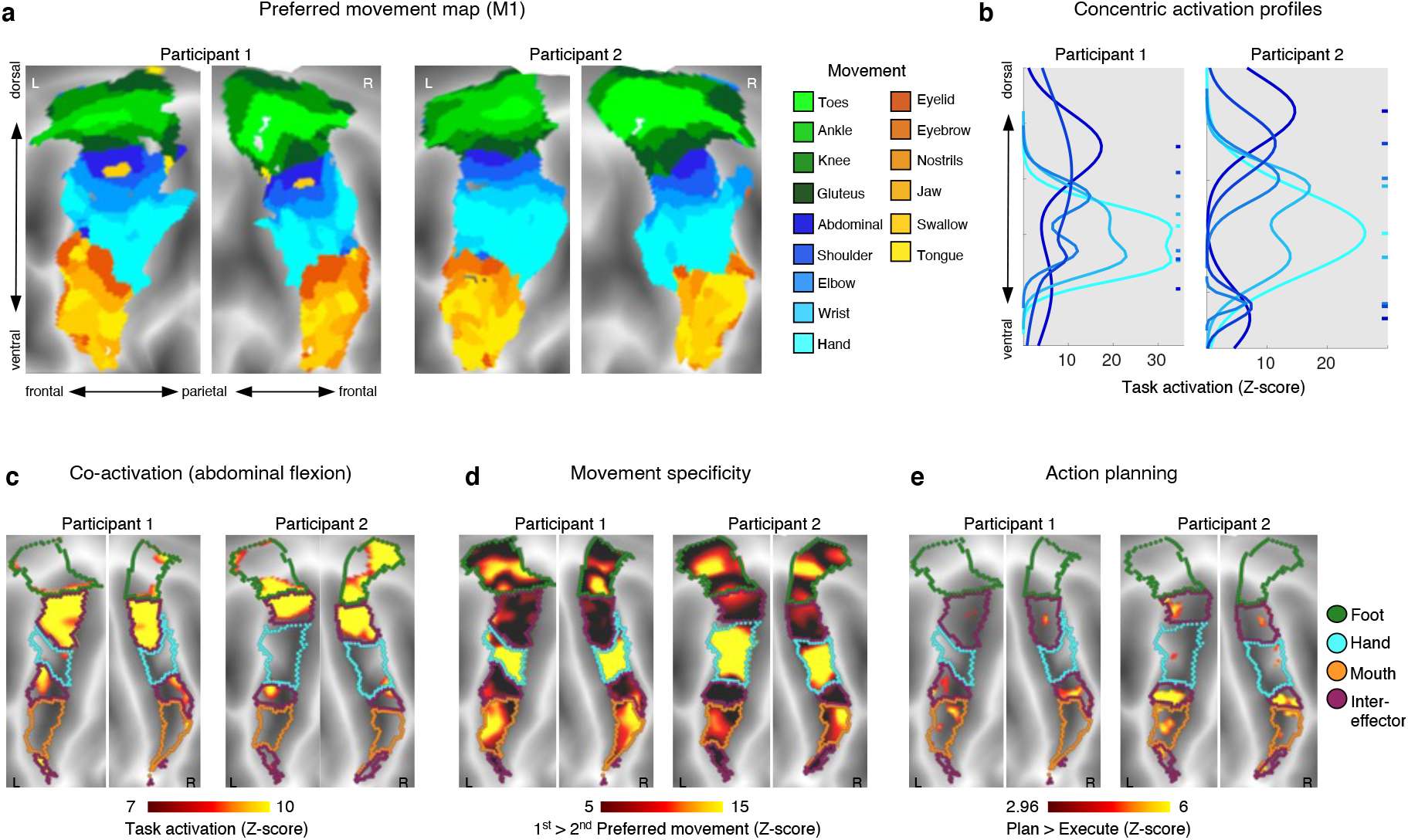
Individual-specific task activations in primary motor cortex. **a**, Task fMRI activations (P1, P2) during a movement task battery, including movement of the toes, ankles, knees, gluteus, abdominals, shoulders, elbows, hands, eyebrows, eyelids, tongue, and swallowing (244 min/participant). Each cortical vertex is colored according to the movement that elicited the strongest task activation (winner-take-all) and is shown on a flattened representation of the cortical surface. Background shading indicates sulcal depths. **b**, Activation strength for each movement was computed along the dorsal-ventral axis within M1. A two-peak gaussian curve was fitted to each movement activation (see Methods). Fitted curves are shown for movement of abdominals, shoulder, elbow, wrist, and hand. Peak locations (hashes on right) were arranged concentrically around the hand peak. See Fig. S7a,b for all movements. **c**, Inter-effector regions were co-activated during abdominal contraction. **d**, Inter-effector regions exhibited more generalized evoked activity during movements. Movement specificity was computed as the activation difference between the first and second-most preferred movements for the six conditions that most activated each discrete region (toes, abdominal, hand, eyelid, tongue, swallowing). **e**, Event-related task fMRI data during an action planning task with separate planning and execution phases for movements of the hands and feet (see Methods). M1 activity in the planning phase was higher than in the execution phase in the inter-effector but not the effector-specific regions.

To formally test for a concentric organization, we fit one- and two-peak gaussian curves to the task activation profiles along the dorsomedial-to-ventrolateral axis of M1. Two-peak curve fits were significantly better for all movements (F-test for comparing models: all *F* > 6.9, all *P* < 0.001) except hand in P2 (F ≅ 0, all *P* ≅ 1) (Fig. S7a). The curve fits revealed concentric activation zones centered around activation peaks for distal movements (hand (Fig. 3b), toes, and tongue (Fig. S7b)) and expanding outward to more proximal movements (shoulder, gluteus, jaw). Concentric rings of activation from separate foot/hand/mouth centers intersected in the top and middle inter-effector regions.

Some movements requiring less fine motor control, such as isometric contraction of the abdominals (Fig. 3c), or raising the eyebrow, co-activated multiple inter-effector regions and the CON (Fig. S8a,b,e). In contrast, movements of the foot and hand only activated the corresponding effector-specific regions (Fig. S8c,d,e). Unlike effector-specific regions, the inter-effectors exhibited only weak movement-specificity, with minimal activation differences between their 1^st^ and 2^nd^ most preferred movements (Fig. 3d).

Regions in CON instantiate action plans, suggesting the CON-to-inter-effector connection could carry general action planning signals. Across foot and hand movements in the novel coordination task, the inter-effectors showed greater activity during action planning than movement execution, but the effector-specific regions did not (Fig. 3e), suggesting that the implementation of action plans may be enabled in part by the inter-effector regions in M1.

Below, we discuss the implications of our findings.

### Similarities between human inter-effector motif and non-human primate anterior M1

When we examined PFM functional connectivity data from a macaque (Fig. S9a), seeds in the posterior bank of precentral gyrus M1 revealed foot, hand, and mouth effector-specific functional connectivity patterns consistent with those seen in humans^29^. Anterior precentral gyrus regions revealed functional connectivity with each other and rostral cingulate zone, pre-SMA, and parietal cortex that appeared analogous to the human CON (Fig. S9a).

In macaques, distinct patterns of corticospinal connectivity distinguish anterior from posterior M1^12,13^. Phylogenetically newer, posterior M1 represents the effectors^14^, projects contralaterally, mainly to the cervical and lumbar enlargements of the spinal cord^13^, and contains more projections that synapse directly onto muscle-innervating spinal neurons^12^ for fine motor control. In contrast, older anterior M1 represents both the body^14^ and more complex actions^4,18,44,45^, projects bilaterally throughout the spinal cord^13^, and connects to internal organs such as the adrenal medulla^10^ and stomach^46^. Non-human primate electrophysiological studies reporting action planning signals often record from sites overlapping with the proposed anteriorly located macaque inter-effector and CON analogues^47^. Thus, direct stimulation, electrophysiological recording, structural and functional connectivity data all reveal similarities between macaque anterior motor cortex and the human inter-effector motif (Fig. S9b).

### Distinct systems for effector isolation and integrated action interleave in motor cortex

Penfield conceptualized his direct stimulation findings in M1 as a continuous map of the human body – the homunculus – an organizational principle that has been dominant for almost 100 years (Fig. 4a). Based on novel and extant data, we instead propose a dual-systems, integrate-isolate model of behavioral control, in which effector isolating and whole-organism action implementation regions alternate (Fig. 4b). This model better fits the human imaging data presented here demonstrating contrasting structural, functional and connectivity patterns within M1. The inter-effector patterning emerges in infancy and is preserved even in the presence of substantial perinatal cortical injury (Fig. S2). In the integrate-isolate model, the regions for foot/hand/mouth fine motor skill are organized somatotopically as three concentric functional zones with distal parts of the effector (toes, fingers, tongue) at the center and proximal ones (knee, shoulder, jaw) on the perimeter. The inter-effector regions at the edges of the effector zones coordinate with each other and the CON to accomplish holistic, whole-body functions in the service of performing actions. The present work suggests these functions include action implementation, as well as postural and gross motor control of axial muscles, while prior work in humans and non-human primates suggests these circuits may also regulate arousal^7^ and control of internal processes and organs (i.e., blood pressure^6^, stomach^46^, adrenal medulla^10^), consistent with circuits for whole-body, metabolic, and physiological control. Thus, the inter-effector system fulfills the role of a Mind-Body Interface (MBI). The MBI forms part of a unified action control system, in conjunction with the CON’s upstream executive control operations, to coordinate gross movements and muscle groups (e.g., torso, eyebrow), and enact top-down control of posture and internal physiology, while preparing for actions. These proposed functions converge with the concept of allostatic regulation, by which the brain anticipates upcoming changes in physiological demands based on planned actions and exerts top-down preparatory control over the body^48^.

**Fig 4.**
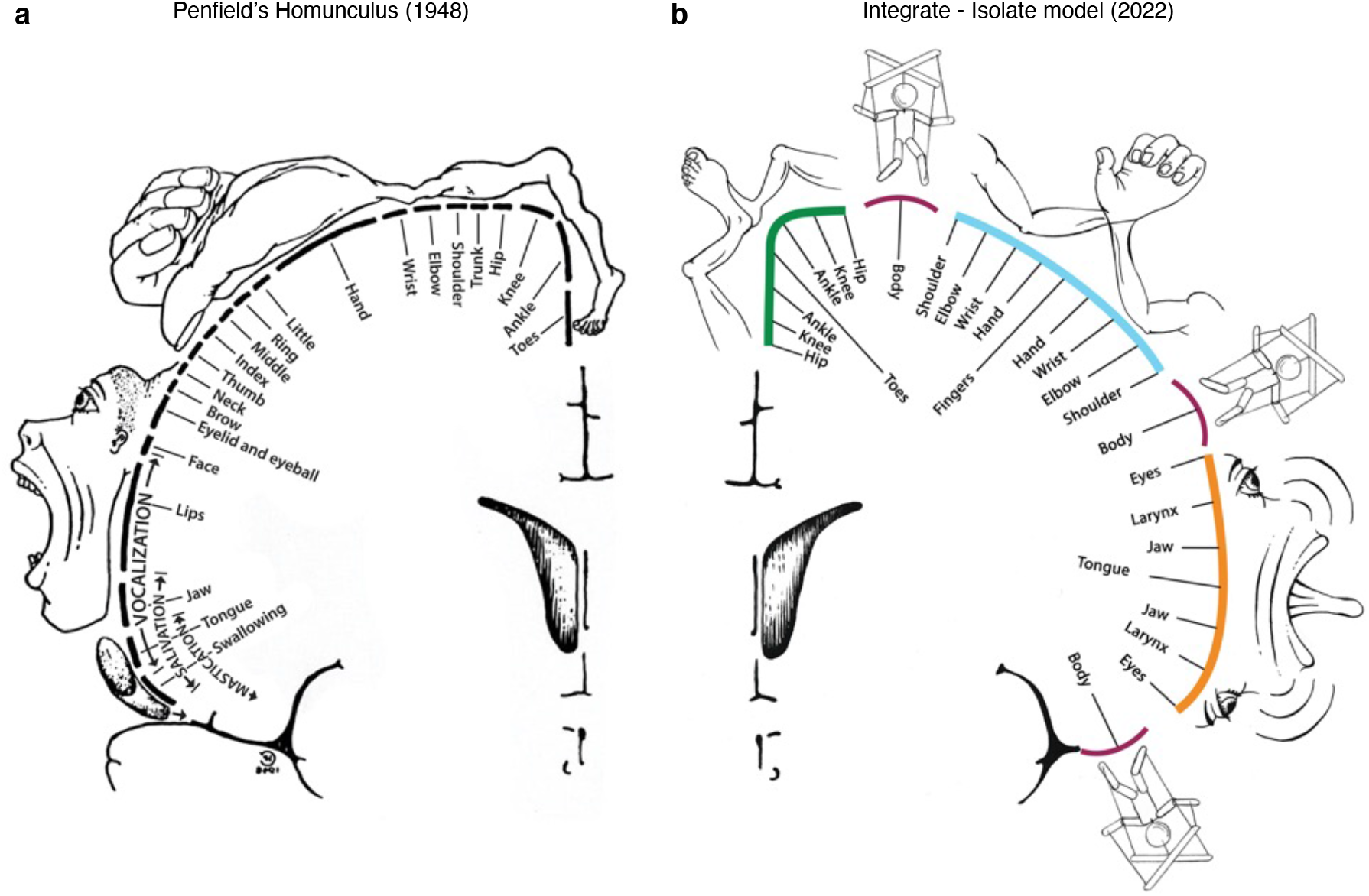
The interrupted homunculus – integrate/isolate model of behavioral and motor control. **a**, Penfield’s classical homunculus (adapted from ^2^) depicting a continuous map of the body in primary motor cortex. **b**, In the integrate-isolate model of primary motor cortex organization, effector-specific (foot [green], hand [cyan], mouth [orange]) functional zones are represented by concentric rings with proximal body parts surrounding the relatively more isolatable distal ones (toes, fingers, tongue). Inter-effector regions (maroon) sit at the intersecting points of these fields, forming part of a Mind-Body Interface (MBI) for integrative, allostatic whole-body control.

### Human stimulation evidence for the integrate-isolate patterning of motor cortex

Penfield proposed the homunculus as an approximation of group-averaged, intraoperative direct electrocortical stimulation data, which showed significant overlap across patients and body parts. He later described his artistic rendering of the homunculus as “an aid to memory […] a cartoon of representation in which scientific accuracy is impossible”^2^. Re-examination of extant human stimulation data raises doubts about the veracity of the homunculus in individuals^11^ and reveals an equal or better fit with the integrate-isolate model. In some patients, a distal-to-proximal concentric organization was documented for the upper limb, just as in non-human primates^49^, while face movements could be elicited from areas dorsal to the hand representation^50^. In addition to focal movements, several other response types are routinely elicited with M1 stimulation, all of which can be better accounted for by whole-organism control regions. Patients have reported the urge to move, while being aware that they are holding still; they have reported a sense of moving though no movement is detectable; or they have moved but denied having done so^2^—effects consistent with modulation of a system also representing action goals. These responses are similar to those typically elicited in CON regions such as dACC^51^ and anterior parietal cortex^52^.

Stimulations almost never produce isolated torso or shoulder movements^49^, and a common outcome of stimulation is no reported response at all^2^. Historically, stimulations failing to elicit movement were not documented. However, we re-analyzed motor stimulations from a recent large study^53^ by mapping them onto cortex, revealing a region that never elicited movement in any patient, corresponding to the middle inter-effector region (Fig. S10a). These results suggest that stimulation strengths deemed safe in humans may not typically elicit movements in the M1 Mind-Body Interface regions, akin to higher-order lateral and medial premotor (i.e., pre-SMA) regions^54^. Human brain-computer-interface (BCI) recordings in M1 have also demonstrated whole-body movement tuning^55^, possibly reflecting inter-effector activity and suggesting that the inter-effector motif could provide a target for whole-body BCI.

### Clinical evidence for separate movement isolation and integrated action control systems

Brain lesion data further support the existence of dual systems for movement isolation and action integration, with partial redundancy in M1. Motor deficits after middle cerebral artery strokes are unilateral, more severe in most distal effectors, and without significant global organismal control deficits^56^. By contrast, lesions of MBI-linked CON regions (dACC, anterior insula, aPFC) can cause isolated volitional deficits ranging from decreased fluency to abulia to akinetic mutism, with preserved motor abilities but little self-generated movement^57,58^. Similarly, anterior motor lesions in macaques can spare visually guided movements while selectively disrupting internally generated actions^59^. Animals with lesions in effector M1 typically recover gross effector control very quickly^60^, while fine finger movement deficits persist longer^61^. More rapid recovery of gross motor abilities may be in part due to proximal functions being taken up by the contra-lesional MBI circuits, enabled by their bilateral spinal cord connectivity. Persistent deficits may therefore be more likely in functions uniquely supported by the effector-specific circuitry.

In a proof-of-principle patient with extensive bilateral perinatal strokes but typical motor ability, extensive post-stroke reorganization maintained the MBI patterning at the cost of sacrificing part of the already reduced M1 hand area^35^. The top third of M1 was destroyed, and surviving cortex contained an M1 hand area that was ventrally shifted and much smaller than in typical controls. Surprisingly, MBI regions were identified both above and below the surviving effector-specific hand region (Fig. S2f), highlighting the Mind-Body Interface’s importance for typical motor ability.

With specific connections to thalamic motor nuclei used as targets for clinical intervention (VIM, CM), the CON-linked MBI may be relevant for a variety of movement disorders, including dystonia or essential tremor (see Supplemental Discussion). Parkinson’s Disease (PD) is of particular note. Many PD symptoms cut across motor, physiological and volitional domains (e.g., postural instability, autonomic dysfunction, and reduced self-initiated activity, among many others^62^), mirroring MBI connections to regions relevant for postural control (cerebellar vermis), volition, and physiological regulation (CON^6,7,57,58^).

### Sensory and motor systems share common functional organization features

Many of the motor cortex organizational features we describe here have clear parallels in sensory systems. Like the concentric somatotopic organization with fine finger movements at the center, primary visual cortex over-represents higher acuity processing at the center, concentrically transitioning to lower acuity in the periphery^63^. Like our integrate/isolate dual-systems model, visual processing streams are parallel and separated in thalamus, early visual cortex, and higher order visual processing streams, with each level of processing maintaining segregation of different types of information (e.g., early: eccentricity vs. angle^64^; late: faces vs. objects^65^). Auditory processing may have similar features, as acoustic signals are processed at least partially in parallel for hearing and speech perception in superior temporal gyrus^66^. These findings suggest shared organizational principles across the brain’s input and output processing streams. It is possible S1 may also have some concentric organizational elements, which should be explored in future work.

### A network for mind-body integration

Two behavioral control systems are interleaved in human motor cortex. One well-known system consists of effector-specific circuits for precise, isolated movements of highly specialized appendages—fingers, toes, and tongue—the type of dexterous motion needed for speaking or manipulating objects. A second, integrative output system, the Mind-Body Interface (MBI) is more important for controlling the organism as a whole. The MBI integrates body control (motor and autonomic) and action planning, consistent with the idea that aspects of higher-level executive control might derive from movement coordination^67^. The MBI includes specific regions of M1, the SMA, thalamus (VIM, CM), posterior putamen, and the postural cerebellum and is functionally connected to the dACC, which has been linked to free will^58^, parietal regions representing movement intentions^52^, insular regions for processing somatosensory, pain^9^, and interoceptive visceral signals^38^. A common factor across this fairly wide range of processes is that they must be integrated if an organism is to achieve its goals through movement while avoiding injury and maintaining physiological allostasis^48^. The MBI provides a substrate for this integration, enabling pre-action anticipatory postural, cardiovascular, and arousal changes (e.g., shoulder tension, increased heart rate, butterflies in the stomach). The discovery that action and body control are melded in a common circuit could help explain why mind and body states so often interact.

## Acknowledgements

The authors would like to thank Dr. Peter Strick for extensive comments and discussions that allowed critical conceptualization of results, as well as Dr. Michael Graziano for helpful comments and suggestions.

This work was supported by NIH grants NS110332 (DJN), MH120989 (CJL), MH100019 (NAS), MH129493 (DMB), MH113883 (CER), EB031765 (JZ), EB031765 (JZ), DA048742 (JZ), MH120194 (JTW), NS123345 (BPK), NS098482 (BPK), NS124789 (SAN), MH118370 (CG), MH122389 (CMS), DA047851 (CL), MH118388 (CL), MH114976 (CL), DA041148 (DAF), DA04112 (DAF), MH115357 (DAF), MH096773 (DAF and NUFD), MH122066 (EMG, DAF, and NUFD), MH121276 (EMG, DAF, and NUFD), MH124567 (EMG, DAF, and NUFD), NS129521 (EMG, DAF, and NUFD), and NS088590 (NUFD); by NSF grant CAREER BCS-2048066 (CG); by the Dystonia Medical Research Foundation (SAN); by the National Spasmodic Dysphonia Association (EMG and SAN); by the Taylor Family Foundation (CMS); by the Intellectual and Developmental Disabilities Research Center (DJG); and by Mallinckrodt Institute of Radiology pilot funding (DJG, EMG).

## METHODS

### Participants

#### Individual-specific Precision Functional Mapping (PFM) data – Washington University adults

##### Participants

Data were collected from three healthy, right-handed, adult participants (ages 35, 25, and 27; 1 female) as part of a study investigating effects of arm immobilization on brain plasticity (data previously published in ^32,33,68^). Two of the participants are authors (NUFD and ANN). Informed consent was obtained from all participants. The study was approved by the Washington University School of Medicine Human Studies Committee and Institutional Review Board. The primary data employed here was collected either prior to the immobilization intervention (Participants 1, 3) or two years afterwards (Participant 2). Data collected immediately after the intervention is presented for within-participant replication in Fig S1b. For details concerning data acquisition and processing, see ^32^.

In two participants (Participants 1, 2), we collected additional fMRI data using the same sequence during performance of two motor tasks: a somatotopic mapping task and a motor control task.

##### Movement task battery

A block design was adapted from the motor task from ^34^. In each run, the participant was presented with visual cues that directed them to perform one of five specific movements. Each block started with a 2.2 s cue indicating which movement was to be made. After this cue, a centrally-presented caret replaced the instruction and flickered once every 1.1 s (without temporal jittering). Each time the caret flickered, participants executed the proper movement. 12 movements were made per block. Each block lasted 15.4 seconds, and each task run consisted of 2 blocks of each type of movement as well as 3 blocks of resting fixation. Movements conducted within each run were as follows:

| Run Type 1 | Run Type 2 | Run Type 3 | Run Type 4 | Run Type 5 | # of runs<br>(P1 / P2) |
| --- | --- | --- | --- | --- | --- |
| Close L hand | Close R hand | Flex L Foot | Flex R Foot | Move<br>Tongue L/R | 24 / 20 |
| Flex L elbow | Flex R Elbow | Flex L Wrist | Flex R Wrist | Lift bilat<br>Shoulders | 10 / 11 |
| Flex L<br>Gluteus | Flex R<br>Gluteus | Tense<br>Abdomen | Open/Close<br>Mouth | Swallow | 10 / 11 |
| Flex L Ankle | Flex R Ankle | Bend L Knee | Bend R Knee | Flex bilat<br>Toes | 10 / 11 |
| Lift L<br>Eyebrow | Lift R<br>Eyebrow | Wink L Eyelid | Wink R<br>Eyelid | Flare Nostrils | 10 / 11 |

##### Action control and coordination task

An event-related design implemented using JSpsych toolbox v6.3 was used to discriminate planning and execution of limb movement. Within the run, the participant is prompted to move either a single limb or to simultaneously move two limbs. There are four possible motions— open-close of fingers or toes, left-right flexion of the wrist/ankle, clockwise rotation of the wrist/ankle, and counterclockwise rotation of the wrist/ankle—each of which may be executed by any of the four extremities (left or right upper/lower extremity). Each motion/extremity combination may be required in isolation, or in combination with a second simultaneous motion. The participant is cued to prepare the movement(s) when they see one or two movement symbols placed on a body shape in a grey color (planning phase), and is then cued to execute the movement(s) when the grey symbol or symbols turn green (execution phase). Using a pseudorandom jitter, the planning phase can last from 2 to 6.5s followed by 4 to 8.5s of movement execution. Each movement trial (planning and execution) is followed by a jittered fixation of up to 5s. A rest block of 8.6s is implemented every 12 movements. Two possible movements are requested during the task run and practiced before the task. The movement pair is changed for each task run. Twelve total runs were acquired per participant.

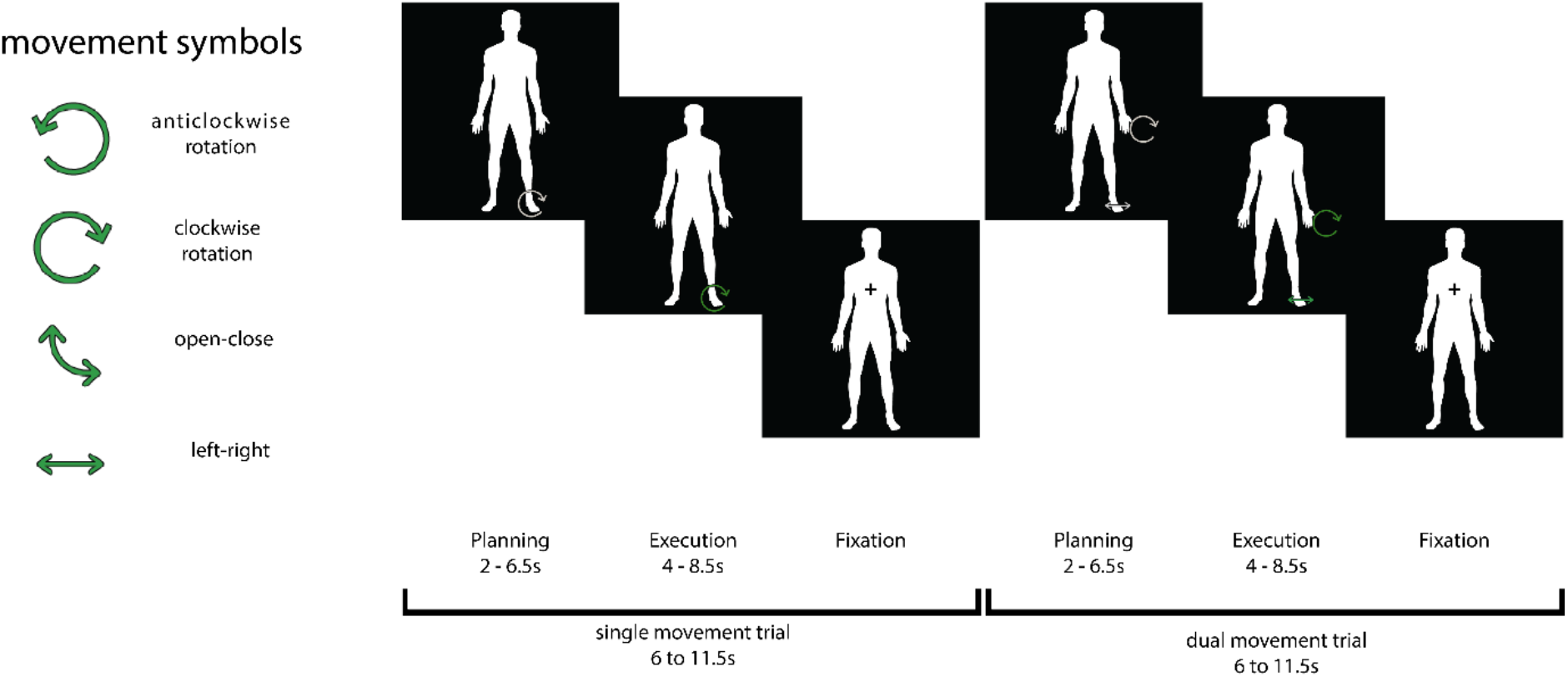

#### Individual-specific PFM data – Cornell adults

##### Participants

Data were collected from four healthy adult participants (ages 29, 38, 24, and 31; all male) as part of a previously published study ^69^. Two of the participants are authors (CJL and JDP). The study was approved by the Weill Cornell Medicine Institutional Review Board.

For details concerning data acquisition and processing, see ^69^.

#### Individual-specific PFM data – Neonate

##### Participant

Data were collected from one sleeping, healthy full-term neonatal participant beginning 13 days after birth, corresponding to 42 weeks post-menstrual age. The study was approved by the Washington University School of Medicine Human Studies Committee and Institutional Review Board.

##### MRI acquisition

The participant was scanned while asleep over the course of 4 consecutive days using a Siemens Prisma 3T scanner on the Washington University Medical Campus. Every session included collection of a high-resolution T2-weighted spin-echo image (TE=563ms, TR=3200ms, flip angle=120°, 208 slices with 0.8×0.8×0.8mm voxels). In each session, a number of 6 minute 45 second multi-echo resting-state fMRI runs were collected as a five-echo blood oxygen level-dependent (BOLD) contrast sensitive gradient echo-planar sequence (flip angle=68°, resolution=2.0 mm isotropic, TR=1761ms, multiband 6 acceleration, TE_1_: 14.20 ms, TE_2_: 38.93 ms, TE_3_: 63.66 ms, TE_4_: 88.39 ms, and TE_5_: 113.12 ms). The number of BOLD runs collected in each session depended on the ability of the neonate to stay asleep during that scan; across the four days, 23 runs were collected in total. A pair of spin echo EPI images with opposite phase encoding directions (AP and PA) but identical geometrical parameters and echo spacing were acquired between every 3 BOLD runs or any time the participant was removed from the scanner.

##### MRI processing

Structural and functional processing followed the pipeline used for the Wash U dataset, with two exceptions. First, segmentation, surface delineation, and atlas registration were conducted using a T2-weighted image (the single highest quality T2 image, as assessed via visual inspection) rather than a T1-weighted image, due to the inverted image contrast observed in neonates. Second, after the multi-echo BOLD data was unwarped and normalized to atlas space, it was optimally combined before nuisance regression and mapping to cifti space. All fMRI scans from the second day of scanning were excluded due to registration abnormalities.

#### Individual-specific PFM data – Infant

##### Participant

Data were collected from one healthy sleeping infant age 11 months. The study was approved by the Washington University School of Medicine Human Studies Committee and Institutional Review Board.

##### MRI acquisition

The participant was scanned while asleep over the course of three sessions using a Siemens Prisma 3T scanner on the Washington University Medical Campus. The first session included collection of a high-resolution T1-weighted MP-RAGE (TE=2.24ms, TR=2400ms, flip angle=8°, 208 slices with 0.8×0.8×0.8mm voxels) and a T2-weighted spin-echo image (TE=564ms, TR=3200ms, flip angle=120°, 208 slices with 0.8×0.8×0.8mm voxels). The second and third sessions included collection of 26 total runs of resting-state fMRI, each collected as a 6 minute 49 second long blood oxygen level-dependent (BOLD) contrast sensitive gradient echo-planar sequence (flip angle=52°, resolution=3.0 mm isotropic, TE=30ms, TR=861ms, multiband 4 acceleration). For each run, a pair of spin echo EPI images with opposite phase encoding directions (AP and PA) but identical geometrical parameters and echo spacing were acquired to correct spatial distortions.

##### MRI processing

Structural processing followed the DCAN Labs processing pipeline (https://github.com/DCAN-Labs/abcd-hcp-pipeline)^70^, which we found performed the best surface segmentation at this age. Functional processing followed the pipeline used for the Washington University adult dataset.

#### Individual-specific PFM data – Child

##### Participant

Data were collected from one healthy awake male child age 9 yo. The study was approved by the Washington University School of Medicine Human Studies Committee and Institutional Review Board.

##### MRI acquisition

The participant was scanned repeatedly over the course of 12 sessions using a Siemens Prisma 3T scanner on the Washington University Medical Campus. These sessions included collection of 14 high-resolution T1-weighted MP-RAGE images (TE=2.90ms, TR=2500ms, flip angle=8°, 176 slices with 1mm isotropic voxels), 14 T2-weighted spin-echo images (TE=564ms, TR=3200ms, flip angle=120°, 176 slices with 1mm isotropic voxels), and 26 total runs of restingstate fMRI, each collected as a 10 minute long blood oxygen level-dependent (BOLD) contrast sensitive gradient echo-planar sequence (flip angle=84°, resolution=2.6mm isotropic, 56 slices, TE=33ms, TR=1100ms, multiband 4 acceleration). In each session, a pair of spin echo EPI images with opposite phase encoding directions (AP and PA) but identical geometrical parameters and echo spacing were acquired to correct spatial distortions in the BOLD data.

##### MRI processing

Structural and functional processing followed the DCAN Labs processing pipeline (https://github.com/DCAN-Labs/abcd-hcp-pipeline) ^70^.

#### Individual-specific PFM data – Perinatal stroke

##### Participant

PS1, a left-handed, 13-year-old male who played for a competitive youth baseball team, was referred to an orthopedic physician because of difficulty using his right arm effectively. Ulnar neuropathy was considered and he was referred for physical therapy. However, PS1 was first seen by a child neurologist (NUFD) for further evaluation. Structural brain MRI revealed unexpectedly extensive bilateral cystic lesions consistent with perinatal infarcts. Review of PS1’s medical history revealed that the injury occurred in the perinatal period.

Data acquisition from PS1 were performed with the approval of the Washington University Institutional Review Board. Written informed consent was provided by PS1’s mother and assent was given by PS1 at the time of data acquisition.

For additional details regarding clinical history, neuropsychological evaluations, motor assessments, or MR image acquisition or processing, see ^35^.

#### Individual-specific PFM data – Macaque

##### Animal Preparation

Data were collected from a sedated adult female macaque monkey (Macaca fascicularis).

Experimental procedures were carried out in accordance with the University of Minnesota Institutional Animal Care and Use Committee and the National Institute of Health standards for the care and use of non-human primates. The subject was fed ad libitum and pair-housed within a light and temperature-controlled colony room. The animal was not water restricted. The subject did not have any prior implant or cranial surgery. The animal was fasted for 14–16 hr prior to imaging. On scanning days, anesthesia was first induced by intramuscular injection of atropine (0.5 mg/kg), ketamine hydrochloride (7.5 mg/kg), and dexmedetomidine (13 μg/kg).

The subject was transported to the scanner anteroom and intubated using an endotracheal tube. Initial anesthesia was maintained using 1.0%–2% isoflurane mixed with oxygen (1 L/m during intubation and 2 L/m during scanning to compensate for the 12-m length of the tubing used). For functional imaging, the isoflurane level was lowered to 1%. The subject was placed onto a custom-built coil bed with integrated head fixation by placing stereo-tactic ear bars into the ear canals. The position of the animal corresponds to the sphinx position. Experiments were performed with the animal freely breathing. Continuous administration of 4.5 μg/kg/hr dexmedetomidine using a syringe pump was administered during the procedure. Rectal temperature (∼99.6F), respiration (10–15 breaths/min), end-tidal CO 2 (25–40), electrocardiogram (70–150 bpm), and SpO2 (>90%) were monitored using an MRI compatible monitor (IRAD-IMED 3880 MRI Monitor, USA). Temperature was maintained using a circulating water bath as well as chemical heating pads and padding for thermal insulation.

##### MRI acquisition

Data were acquired on a Siemens Magnetom 10.5 T Plus. A custom in-house built and designed RF coil was used with an 8-channel transmit/receive end-loaded dipole array of 18-cm length combined with a close-fitting 16-channel loop receive array head cap, and an 8-channel loop receive array of 50×100 mm2 size located under the chin^71^. A B1+ (transmit B1) fieldmap was acquired using a vendor provided flip angle mapping sequence and then power calibrated for each individual. Following B1+ transmit calibration, 3–5 averages (23 min) of a T1 weighted MP-RAGE were acquired for anatomical processing (TR = 3300 ms, TE = 3.56 ms, TI = 1140, flip angle = 5°, slices = 256, matrix = 320×260, acquisition voxel size = 0.5×0.5×0.5 mm 3, inplane acceleration GRAPPA = 2). A resolution and FOV matched T2 weighted 3D turbo spin-echo sequence was run to facilitate B1 inhomogeneity correction. Five images were acquired in both phase-encoding directions (R/L and L/R) for offline EPI distortion correction. Six runs of fMRI timeseries, each consisting of 700 continuous 2D multiband EPI^72–74^ functional volumes (TR = 1110ms; TE = 17.6 ms; flip angle = 60°, slices = 58, matrix = 108×154; FOV = 81×115.5mm ; acquisition voxel size = 0.75×0.75×0.75 mm) were acquired with a left-right phase encoding direction using in plane acceleration factor GRAPPA = 3, partial Fourier = 7/8th, and MB factor = 2. Since the macaque was scanned in sphinx position, the orientations noted here are what is consistent with a (head first supine) typical human brain study (in terms of gradients) but translate differently to the actual macaque orientation.

##### MRI processing

Processing followed the DCAN Labs non-human primate processing pipeline (https://github.com/DCAN-Labs/nhp-abcd-bids-pipeline), with minor modifications. Specifically, we observed that distortion from the 10T scanner was so extensive that the field maps did not fully correct it. Therefore, instead of field-map based unwarping, we used the computed field map-based warp as an initial starting point for Synth, a field map-less distortion correction algorithm that creates synthetic undistorted BOLD images for registration to anatomical images^75^. Synth substantially reduced residual BOLD image distortion.

#### Group-averaged datasets

Resting-state fMRI data was averaged across participants within each of five large datasets. <u>UK Biobank (UKB):</u>

A group-average weighted eigenvectors file from an initial batch of 4100 UKB participants scanned using resting-state fMRI for six minutes was downloaded from https://www.fmrib.ox.ac.uk/ukbiobank/. This file consisted of the top 1200 weighted spatial eigenvectors from a group-averaged PCA. See ^76^ and documentation at https://biobank.ctsu.ox.ac.uk/crystal/ukb/docs/brain_mri.pdf for details of the acquisition and processing pipeline. This eigenvectors file was mapped to the Conte69 surface template atlas^77^ using the ribbon-constrained method in Connectome Workbench^78^, and the eigenvector timecourses of all surface vertices were cross-correlated.

##### Adolescent Brain Cognitive Development (ABCD) Study

Twenty minutes (4 × 5 minute runs) of resting-state fMRI data, as well as high-resolution T1-weighted and T2-weighted images, were collected from 3,928 9-10 year old participants, who were selected as the participants with at least 8 minutes of low-motion data from a larger scanning sample. Data collection was performed across 21 sites within the Unites States, harmonized across Siemens, Philips, and GE 3T MRI scanners. See^79^ for details of the acquisition parameters. Data processing was conducted using the ABCD-BIDS pipeline (https://github.com/DCAN-Labs/abcd-hcp-pipelines); see^80^ for details.

##### Human Connectome Project (HCP)

A vertexwise group-averaged functional connectivity matrix from the HCP 1200 participants release was downloaded from db.humanconnectome.org. This matrix consisted of the average strength of functional connectivity across all 812 participants who completed four 15-minute resting-state fMRI runs and who had their raw data reconstructed using the newer “recon 2” software. See ^78,81–83^ for details of the acquisition and processing pipeline.

##### Washington University 120 (WashU 120)

Data was collected from 120 healthy young adult participants recruited from the Washington University community during relaxed eyes-open fixation (60 females, mean age = 25 years, age range = 19-32 years). Scanning was conducted using a Siemens TRIO 3.0T scanner and included collection of high-resolution T1-weighted and T2-weighted images, as well as an average of 14 minutes of resting-state fMRI. See ^84^ for details of the acquisition and processing pipeline.

##### Neonates (eLABE)

Mothers were recruited during the 2nd or 3rd trimester from two obstetrics clinics at Washington University as part of the Early Life Adversity, Biological Embedding, and Risk for Developmental Precursors of Mental Disorders (eLABE) study. Neuroimaging was conducted in full-term, healthy neonate offspring shortly after birth (average post-menstrual age of included participants 41.4 weeks, range 38-45). Of the 385 participants scanned for eLABE, 262 were included in the current analyses. See ^85^ for additional details of the participants, criteria for exclusion, scanning acquisition protocol and parameters, and processing pipeline.

### Analyses

#### Functional connectivity

For each single-participant dataset, a vertex/voxelwise functional connectivity matrix was calculated from the resting-state fMRI data as the Fisher-transformed pairwise correlation of the timeseries of all vertices/voxels in the brain. In the ABCD, WashU120, and eLABE datasets, vertex/voxelwise group-averaged functional connectivity matrices were constructed by first calculating the vertex/voxelwise functional connectivity within each participant as the Fisher-transformed pairwise correlation of the timeseries of all vertices/voxels in the brain, and then averaging these values across participants at each vertex/voxel.

##### Seed-based functional connectivity

We defined a continuous line of seeds down the left precentral gyrus by selecting every vertex in a continuous straight line on the cortical surface between the most ventral aspect of the medial motor area (approximate MNI coordinates [-4, -31, 54]) and the ventral lip of the precentral gyrus right above the operculum (approximate MNI coordinates [-58 4 8]). For each seed, we examined its map of functional connectivity as the Fisher-transformed correlation between that vertex’s timecourse and that of every other vertex/voxel in the brain.

##### Network detection in somatomotor cortex

To define the somatomotor regions that were visually identified from the seed-based connectivity analysis in an unbiased fashion for further exploration, we entered each individual adult human participant’s data into a data-driven network detection algorithm designed to identify network subdivisions that are hierarchically below the level of classic large-scale networks (e.g. that produce hand/foot divisions in somatomotor cortex; ^27,28^). We have previously described how this approach identifies sub-network structures that converge with task-activated regions^86^ and with known neuroanatomical systems^87^.

In each adult participant, this analysis clearly identified network structures corresponding to motor representation of the foot, hand, and mouth; and it additionally identified network structures corresponding exactly to the previously unknown connectivity pattern identified from the seed-based connectivity exploration as the inter-effector regions. For simplicity, we manually grouped all inter-effector subnetworks together as a single putative network structure (labeled as inter-effector) for further analysis.

Finally, to identify classic large-scale networks in each participant, we repeated the Infomap algorithm on matrices thresholded at a series of denser thresholds (ranging from 0.2% to 5%), and additionally identified individual-specific networks corresponding to the Default, Medial and Lateral Visual, Cingulo-opercular, Fronto-Parietal, Dorsal Attention, Language, Salience, Parietal Memory, and Contextual Association networks following procedures described in ^29^.

##### Differences in functional connectivity between inter-effector and foot/hand/mouth regions

Within each adult human participant, we calculated an inter-effector connectivity map as the Fisher-transformed correlation between the average timecourse of all cortical inter-effector vertices and the timecourse of every other vertex/voxel in the brain. We then repeated this procedure to calculate a connectivity map for the foot, hand, and mouth areas.

To identify brain regions more strongly connected to inter-effector regions than to other motor regions, we computed the smallest positive difference in each voxel/vertex between inter-effector connectivity and any foot/hand/mouth connectivity. That is, we calculated (inter-effector – max[foot, hand, mouth]). This represents a conservative approach that only identifies regions of the brain for which the inter-effector regions are more strongly connected than any of the other motor areas.

##### Functional connectivity with CON

Within each adult human participant, we calculated the functional connectivity between each of the foot, hand, mouth, and inter-effector regions and the CON. This was computed as the Fisher-transformed correlation between 1) the average timecourse across all vertices in the motor region and 2) the average timecourse across all vertices in the CON. We conducted paired t-tests across subjects comparing the inter-effector connectivity with CON against each of the foot/hand/mouth connectivity strengths, correcting for the three tests conducted.

##### Motor/CON network visualization

Visualization of network relationships was conducted using spring-embedded plots ^27^, as implemented in Gephi (https://gephi.org/). In each individual adult human participant, nodes were defined as congruent clusters of foot, hand, mouth, inter-effector, and CON networks larger than 20mm^2^. Pairwise connectivity between nodes was calculated as the Fisher-transformed correlation of their mean timecourses. For visualization purposes, graphs were constructed by thresholding the pairwise node-to-node connectivity matrices at 40% density (the general appearance of the graphs did not change across a range of densities).

##### Functional connectivity with adjacent postcentral gyrus

In each adult human participant, we defined the pre- and postcentral gyri based on the individual-specific Brodmann areal parcellation produced by Freesurfer, which was deformed into fs_LR_32k space to match the functional data. Precentral gyrus was considered to be the vertices labeled as Brodmann Areas 4a and 4p, while postcentral gyrus was the vertices labeled as Brodmann Areas 3b and 2. Brodmann area 3a (fundus of central sulcus) was not considered for this analysis. Because the medial aspect of somatomotor cortex (corresponding to representation of the leg and foot) was always classified by Freesurfer as BA 4a, we defined the medial postcentral gyrus as the cortical vertices with y-coordinates farther posterior than the median y-coordinate of the foot region (from the network mapping above).

Within the participant’s precentral gyrus, we labeled vertices as representing foot, hand, mouth, or inter-effector according to their labels from the network mapping procedure. We then partitioned the postcentral gyrus into foot, hand, mouth, and inter-effector areas depending on which precentral region each vertex was physically closest to. Finally, within each partition (foot, hand, mouth, and inter-effector) we calculated the average connectivity between the pre and postcentral gyrus as the Fisher-transformed correlation between the average timecourses of all vertices in each area. We then conducted paired t-tests across subjects comparing the inter-effector connectivity with adjacent S1 against each of the foot/hand/mouth connectivity strengths with S1, FDR-correcting for the three tests conducted.

##### Functional connectivity with middle insula

In each adult human participant, we defined the middle insula based on the individual-specific Freesurfer gyral parcellation using the Destrieux atlas^88^, which was deformed into fs_LR_32k space to match the functional data. Middle insula was considered to be the vertices labeled as the superior segment of the circular sulcus of the insula or as the short insular gyrus. We then calculated the functional connectivity between each of the bilateral foot, hand, mouth, and inter-effector regions and the bilateral middle insula. We conducted paired t-tests across subjects comparing the inter-effector connectivity with middle insula against each of the foot/hand/mouth connectivity strengths, FDR-correcting for the number of tests conducted.

##### Functional connectivity with cerebellum

In each adult human participant, we calculated the functional connectivity between each of the foot, hand, mouth, and inter-effector regions with each voxel of the cerebellum. Cerebellar connectivity strengths calculated this way were then mapped onto a cerebellar flat map using the SUIT toolbox^89^. Connectivity strengths were averaged within each of 27 atlas regions ^90^. For each region, we conducted three paired t-tests comparing inter-effector connectivity strength against foot, hand, and mouth connectivity strength, FDR-correcting for the total number of tests conducted. Regions were reported if the inter-effector connectivity strength was significantly higher than the connectivity strength of all other motor regions.

##### Functional connectivity with putamen

In each adult human participant, we divided each unilateral putamen in each hemisphere into quarters by splitting it based on the median of its Y (anterior-posterior) and Z (dorsalventral) coordinates. We then calculated the functional connectivity between each of the foot, hand, mouth, and inter-effector regions and each putamen quarter.

For each putamen division, we conducted paired t-tests across subjects comparing the inter-effector connectivity with that putamen division against each of the foot/hand/mouth connectivity strengths, FDR-correcting for the number of tests conducted. We reported divisions in which the inter-effector connectivity was significantly different from all three effector-specific connectivities.

##### Functional connectivity with thalamus

To investigate subregions of thalamus, we employed the DISTAL atlas v1.1 ^91^, which contains a number of histological thalamic subregions identified by ^92^. This atlas was downsampled into the 2mm isotropic space of the functional data. Functional connectivity maps seeded from the foot, hand, mouth, and inter-effector regions in each adult human participant were computed, and mean connectivity values were calculated within each atlas region. The atlas specifies multiple subregions for many nuclei; these subregions were combined and treated as single nuclei for the purposes of connectivity calculation.

For each adult human participant, we averaged the connectivity seeded from the inter-effector regions and from each of the foot, hand, and mouth regions across all voxels within each thalamic nucleus. For each thalamic nucleus, we conducted paired t-tests across subjects comparing the inter-effector with the mean of the foot/hand/mouth connectivity strengths, bonferroni correcting for the number of thalamic nuclei tested.

##### Lag structure of RSFC

We used a previously published method for estimating relative time delays (“lags”) in fMRI data ^36,93^. Briefly, for each session in each adult human participant, we computed a lagged cross-covariance function (CCF) between each pair of vertex/voxel timecourses within the motor system and CON in the cortex. Lags were more precisely determined by estimating the cross-covariance extremum of the session-level CCF using three-point parabolic interpolation. The resulting set of lags was assembled into an antisymmetric matrix capturing all possible pairwise time delays (TD matrix) for each session, which was averaged across sessions to yield participant-level TD matrices. Finally, each participant’s TD matrix was averaged across rows to summarize the average time-shift from one vertex to all other vertices. Average time lag was then averaged across all vertices with each of the precentral gyrus foot/hand/mouth/inter-effector regions, and the CON.

We then conducted paired t-tests across subjects comparing 1) the mean lag in inter-effector regions against the mean lags in each of the foot/hand/mouth regions, and 3) the mean lag in CON regions against the mean lags in each of the foot/hand/mouth regions.

#### Structural MRI

##### Cortical thickness

Within each adult human participant, the map of cortical thickness generated by the Freesurfer segmentation was deformed into fs_LR_32k space to match the functional data. Precentral gyrus foot, hand, mouth, and inter-effector regions were defined as above, and mean cortical thickness was calculated within each region. We then conducted paired t-tests across subjects comparing the inter-effector thickness against each of the foot/hand/mouth thicknesses, correcting for the three tests conducted.

##### Fractional Anisotropy

White matter fibers tracked from separate areas of motor cortex using diffusion imaging quickly converge into the internal capsule and become difficult to dissociate. As such, we tested for FA differences in the white matter immediately below the precentral gyrus.

To calculate FA beneath the cortex, we first constructed fs_LR_32k-space surfaces 2 mm below each gray-white surface in adult human participants P1-P3. To accomplish this, for each vertex on the surface, we computed the 3D vector between corresponding points on the fs_LR_32k pial and the gray-white surfaces, and we extended that vector an additional 2 mm beyond the gray-white surface in order to create a lower surface. We then mapped the FA values using the using the ribbon-constrained method, mapping between the gray-white and the 2mm-under surfaces. The result is FA values mapped to a lower surface within white matter that is in register to the existing fs_LR_32k surfaces on which the functional data is mapped and the motor regions defined. Precentral gyrus foot, hand, mouth, and inter-effector regions were defined as above, and we calculated mean FA beneath each cortical region. We then conducted paired t-tests across subjects comparing the mean FA beneath the inter-effector regions against mean FA beneath each of the foot/hand/mouth thicknesses.

##### Myelin density

Within each adult human participant, we created vertexwise maps of intracortical myelin content following methods described in ^94^ and ^78^. Precentral gyrus was defined as above. Across participants, we found that baseline myelin density values (both in precentral gyrus and in the whole-brain myelin density map) varied wildly across participants in different datasets, likely based on differences in the T1- and T2-weighted sequences employed. Thus, for optimal visualization of results, in each participant we normalized the myelin density values by dividing the calculated vertexwise myelin densities in precentral gyrus by the mean myelin density across the whole precentral gyrus. Finally, precentral gyrus foot, hand, mouth, and inter-effector regions were defined as above, and mean normalized myelin density was calculated within each region. We then conducted paired t-tests across subjects comparing the inter-effector myelin density against each of the foot/hand/mouth myelin densities, correcting for the three tests conducted.

#### Task fMRI

##### Movement task battery analysis

Basic analysis of the movement task battery data was conducted using within-participant block designs. To compute the overall degree of activation in response to each motion, data from each run was entered into a first-level analysis within FSL’s FEAT ^95^ in which each motion block was modeled as an event of duration 15.4s, and the combined block waveform for each motion condition was convolved with a hemodynamic response function to form a separate regressor in a GLM analysis testing for the effect of the multiple condition regressors on the timecourse of activity within every vertex/voxel in the brain. Beta value maps for each condition were extracted for each run and entered into a second-level analysis, in which run-level condition betas were tested against a null hypothesis of zero activation in a one-sample t-test across runs (within participant). The resulting t-values from each motion condition tested in this second-level analysis were converted to Z-scores. Z-score activation maps were smoothed with a geodesic 2D (for surface data) or Euclidean 3D (for volumetric data) Gaussian kernel of σ =2.55 mm.

##### Movement task battery winner take all

For each vertex within the broad central sulcus area, we identified the movement that produced the greatest activation strength (Z-score from second-level analysis, above) in that vertex, and we assigned that motion to that vertex.

##### Movement task battery curve fitting

For each vertex within precentral gyrus, we first computed its position along the dorsal-ventral axis of left hemisphere M1. This was done by identifying the closest point within the continuous line of points running down precentral gyrus (defined in <u>Seed-based functional connectivity)</u>, and assigning that closest point’s ordered position within the line to the vertex.

For every movement, we then plotted that dorsal-ventral M1 position against Z-score activation in each vertex. We then fit two curves to each of these relationships. The first curve was a single-gaussian model of the form:

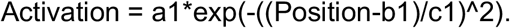

The second curve was a double-gaussian model of the form:

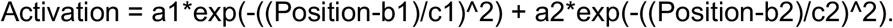

The “a1” and “a2” parameters in each model were constrained to be positive (to enforce positive-going peaks). Curve fitting was constrained to be conducted within the general vicinity of the activated area in order to avoid fitting negative activations observed in distant portions of M1. For lower extremity movements, this meant excluding the bottom third of M1; for upper extremity movements, the bottom third of M1 plus the medial wall; for face movements, the top third of M1.

Finally, we tested whether the one- or two-peak models better fit the data. This was done by conducting an F-test between the models, computed as:

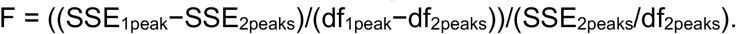

The p value was computed from this F by employing the F-statistic continuous distribution function (fcdf.m) in Matlab and using (df_1peak_ – df_2peaks_) and df_2peaks_ as the numerator and denominator degrees of freedom, respectively.

##### Movement selectivity

Based on results from the above winner-take-all analysis, we identified the movement that was most preferred at the center of each the three effector-specific (toe movement, hand movement, tongue movement) and inter-effector regions (abdominal movement, eyelid movement, swallowing). The center-most movements were selected to avoid issues with spreading, overlapping activation near the borders of effector-specific and inter-effector regions. For every vertex within the precentral gyrus, we compared the strength of activation between the most preferred of the six movements at that vertex against the activation of the second-most preferred movements. The differences between these activation strengths was taken to be the movement selectivity of that vertex.

##### Movement coactivation

For each region among the six resting state-derived foot, hand, mouth, and inter-effector regions in the precentral gyrus, we calculated the average activation within that region for each movement, producing a profile of motor activation strengths for that region. We also calculated the average activation within all CON vertices for each movement. To determine the degree to which various regions were coactive across movements, we then correlated each foot, hand, mouth, and inter-effector cluster’s profile of activation strengths with that of all other clusters, and with that of the CON. Note: visualization of activation maps revealed some striping, suggesting that the Open/Close Mouth and the Bend L Knee conditions were partially distorted by head motion; therefore, these conditions were excluded from analysis, though their inclusion did not change results.

##### Action control and coordination task analyses

Analysis of the action control task was conducted using within-participant event related designs. For each separate run, a GLM model was constructed in FEAT ^95^ in which separate regressors described the initiation of 1) planning and 2) execution of each type of movement (4 movements * 4 limbs). Each regressor was constructed as a 0-length event convolved with a canonical hemodynamic response, and beta values for each regressor were estimated for every voxel in the brain. These beta value maps for each condition were thus computed for each run and entered into a second-level analysis, in which a t-test across runs contrasted the run-level planning betas against the run-level execution betas.

#### Human direct electrocortical stimulation site mapping

Each stimulation location reported by ^53^ was separately mapped into the MNI-space Conte69 atlas pial cortical surface ^77^ by identifying the vertex with the minimal Euclidean distance to the stimulation site’s MNI coordinates. Movements resulting from each site were classified as “lower extremity”, “upper extremity”, or “face” and colored accordingly (though no lower extremity movements were reported in the displayed left hemisphere).

