## Supplemental Material for "A mind-body interface alternates with effector-specific regions in motor cortex"

**SUPPLEMENTARY MATERIALS**

| | P 1 | P 2 | P 3 | P 4 | P 5 | P 6 | P 7 | Avg |
|---|---|---|---|---|---|---|---|---|
| **M1 Cortex** | | | | | | | | |
| L Top | -17 -33 56 | -23 -38 52 | -18 -37 54 | -17 -33 61 | -19 -31 65 | -16 -31 66 | -18 -33 60 | -19 -34 59 |
| R Top | 16 -36 57 | 20 -35 51 | 18 -36 52 | 17 -27 59 | 26 -28 59 | 20 -30 62 | 23 -28 65 | 20 -31 58 |
| L Middle | -39 -20 43 | -36 -21 39 | -39 -23 42 | -41 -15 42 | -36 -15 49 | -38 -15 43 | -38 -15 48 | -38 -18 44 |
| R Middle | 37 -19 49 | 35 -23 39 | 41 -14 44 | 36 -13 42 | 39 -13 38 | 44 -10 48 | 45 -12 42 | 40 -15 43 |
| L Bottom | -54 -5 17 | -55 -12 11 | -55 -2 16 | -51 -3 21 | -55 3 4 | -54 -3 13 | -53 -1 14 | -54 -3 14 |
| R Bottom | 54 -8 16 | 59 -10 12 | 51 0 20 | 62 -1 21 | 57 4 8 | 56 1 15 | 58 4 17 | 56 -1 16 |
| **Putamen** | | | | | | | | |
| L posterior | -35 -4 -4 | -29 -15 3 | -30 -16 3 | -25 0 8 | -25 -1 -10 | -26 -5 13 | -25 -5 -6 | -28 -5 -1 |
| R posterior | 29 -14 6 | 29 -14 1 | 28 -9 1 | 25 -4 8 | 30 -10 6 | 29 -12 10 | 27 2 -9 | 28 -9 3 |
| **Thalamus** | | | | | | | | |
| L centromedian | -9 -23 1 | -10 -22 1 | -11 -22 1 | -9 -20 4 | -13 -22 2 | -10 -20 3 | -10 -20 4 | -10 -21 2 |
| R centromedian | 11 -22 2 | 10 -21 1 | 14 -20 0 | 9 -19 6 | 14 -21 4 | 13 -18 5 | 12 -18 6 | 12 -20 3 |
| **Cerebellum** | | | | | | | | |
| L | -11 -66 -52 | -29 -59 -26 | -34 -53 -53 | -26 -63 -57 | -27 -60 -21 | -11 -66 -45 | -32 -57 -49 | -24 -61 -43 |
| R | 20 -67 -52 | 15 -67 -52 | 9 -67 -12 | 33 -55 -50 | 22 -61 -17 | 28 -47 -48 | 8 -71 -14 | 19 -62 -35 |

**Table S1: Inter-effector regions of interest**
Location of inter-effector regions in each participant, and the average location across participants. Cortical coordinates are centroids of regions; subcortical coordinates are locations of inter-effector functional connectivity peaks within each structure. Coordinates are represented as [X Y Z] in MNI space.

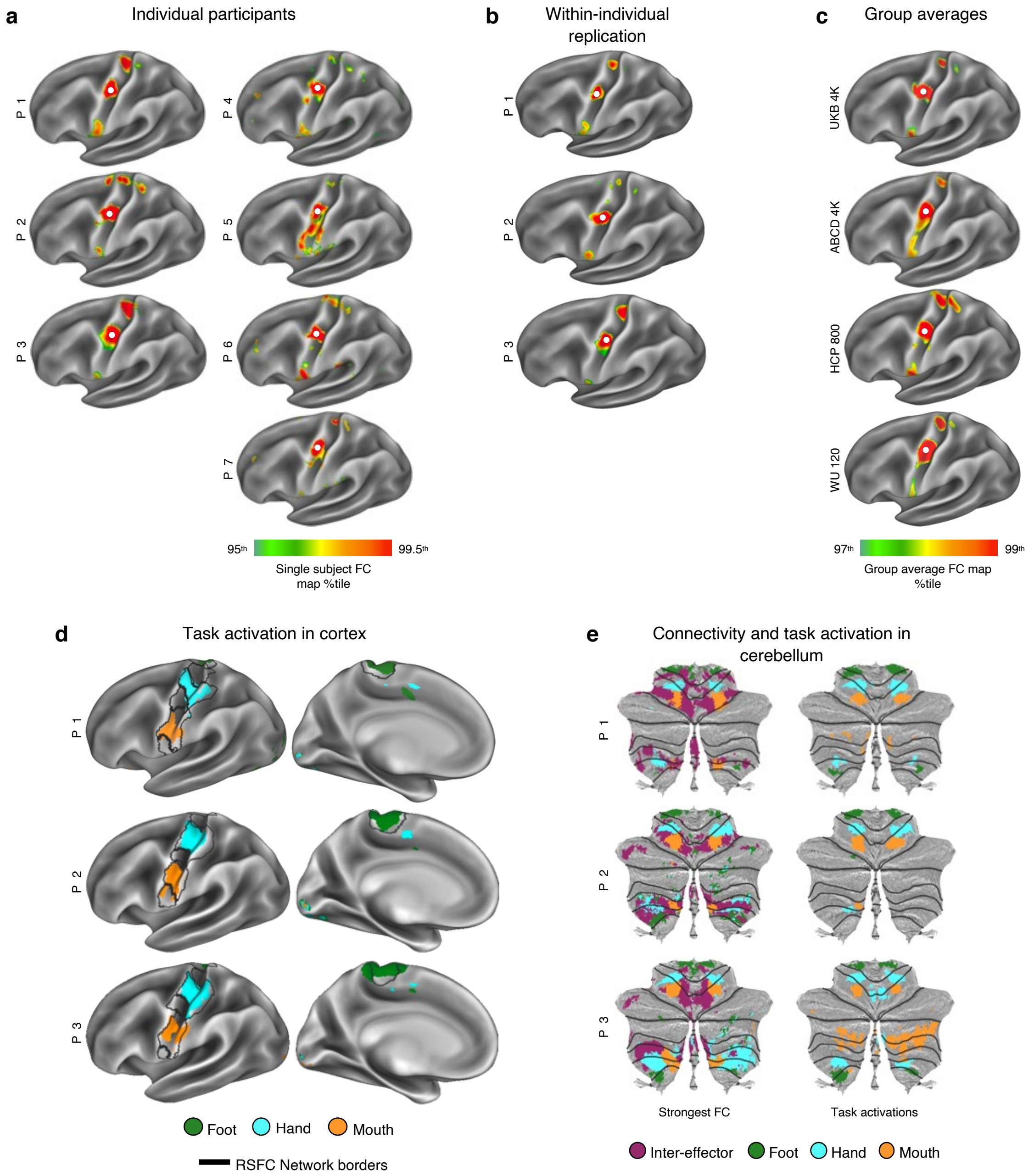


**Fig. S1| Consistency of inter-effector functional connectivity motif.** Connectivity patterns seeded from a continuous line down the left precentral gyrus revealed that the interleaved motor functional connectivity pattern was consistent across **a,** seven highly-sampled individual participants (172-356 minutes of data); **b,** replication data (416-1,114 minutes) collected in P1-P3; and **c,** multiple independent sets of group data averaged across cohorts of varying size. Here, functional connectivity is shown seeded from the middle inter-effector region for each individual participant and group average dataset (see Supplementary Movie 2 for all seeds). Thresholds for connectivity maps were scaled to the 95$^{th}$ percentile of map values in individuals, and to the 97$^{th}$ percentile of values in groups, to account for differences in data acquisition and processing strategies across datasets. **d,** Discrete functional networks were demarcated within each subject in M1 and S1 using a whole-brain, data-driven hierarchical approach applied to the resting-state fMRI data (see Fig S2), which defined the spatial extent of the networks observed in Fig 1 (black outlines). In P1-P3, regions defined by RSFC were functionally labeled using a classic block-design fMRI motor task involving separate movement of the foot, hand, and tongue (following [34]; see [32] for details). The map illustrates the top 1% of vertices activated by movement of the foot (green), hand (cyan), and mouth (orange). **e,** Left: preferential connectivity of each motor division to the cerebellum. Right: activations driven by the fMRI motor task described in panel d. The map illustrates the top 5% of vertices within cerebellum activated by movement of the foot (green), hand (cyan), and mouth (orange).

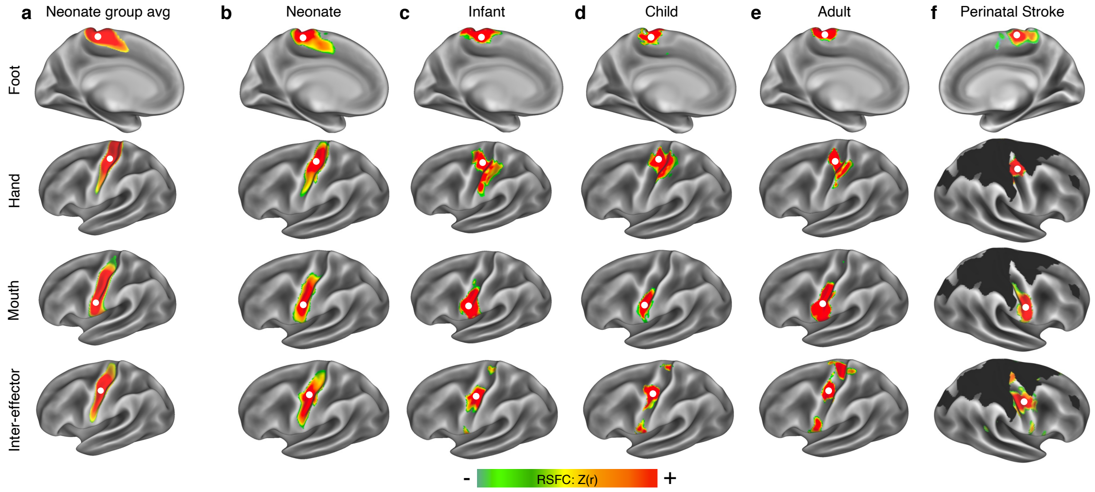


**Fig. S2| Motor cortex functional connectivity in pediatric participants and perinatal stroke.** Functional connectivity maps were seeded from a continuous line of points down precentral gyrus in fMRI data from **a,** data averaged across 262 human neonates, all scanned shortly after birth; **b,** a neonate scanned 13 days after birth; **c,** an 11-month old infant; **d,** a 9-year old child; **e,** adult participant P1 (from Fig. 1); and **f,** an adolescent who had experienced extensive cortical reorganization after severe bilateral perinatal strokes (destroyed cortex in black). Right hemisphere is shown in the stroke patient because left hemisphere M1 was entirely lost. Example seed maps shown here illustrate observed effector-specific (first three rows) and inter-effector (fourth row) connectivity. Effector-specific and inter-effector regions exhibited clear boundaries within M1 in the infant, child, the adults, and the stroke patient, but not in the neonates. Visualization thresholds varied between 0.3 and 0.5 across datasets due to differences in data collection and processing, as well as differences inherent to the populations.

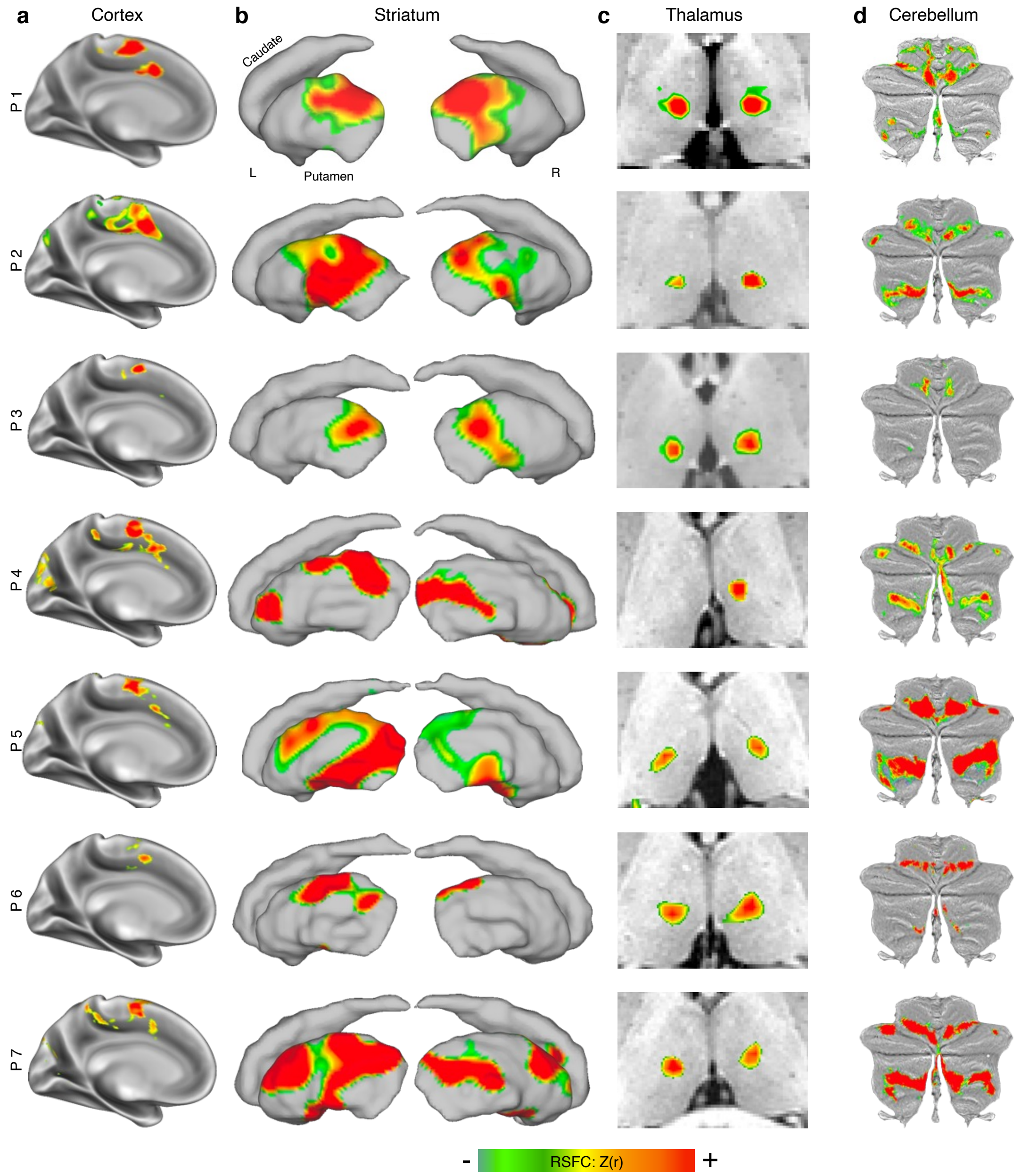


**Fig. S3| Whole brain functional connectivity of inter-effector motif across participants.** Brain regions with the strongest functional connectivity to the middle inter-effector region in **a,** medial cortex, **b,** striatum (lateral view of left and right striatum), **c,** thalamus (axial view), and **d,** cerebellum. Functional connectivity values are thresholded at Z(r) > 0.35 in cortex. Subcortical functional connectivity values are thresholded at different levels in each subject due to variation in subcortical signal-to-noise ratios across individuals. Thresholds were chosen to illustrate the strongest subcortical connections. Specific thresholds shown here: P1 - Z(r) > 0.15; P3, 4, 6, 7 - Z(r) > 0.1; P2 - Z(r) > 0.04; P5 - Z(r) > 0.03.

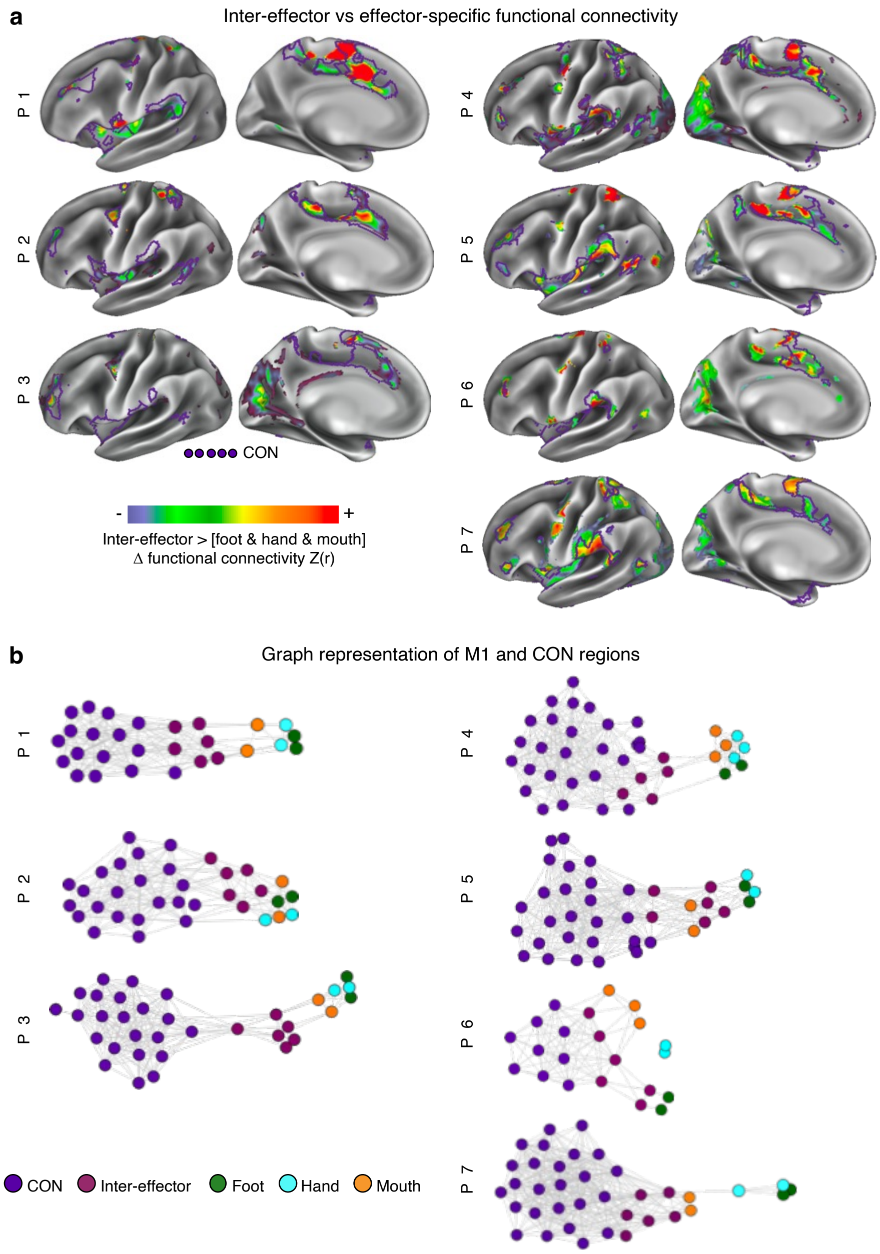


**Fig. S4| Inter-effector regions are strongly connected to CON across subjects. a,** To quantify functional connectivity differences between the foot/hand/mouth effector and inter-effector regions, we created individual-specific connectivity maps seeded from each of the foot/hand/mouth/inter-effector regions. In every subject, we mapped brain regions more strongly functionally connected to the inter-effector motif than any of the foot/hand/mouth regions. The purple outlines show the individual-specific Cingulo-opercular network (CON). Central sulcus regions are masked as they exhibit large differences by definition. **b,** For every participant, relationships between CON, inter-effector, and effector-specific regions are visualized in network space using a spring-embedding plot, in which network nodes are positioned in a 2D plane and connected regions are pulled together while disconnected regions are pushed apart. Connecting lines indicate a strong functional connection.

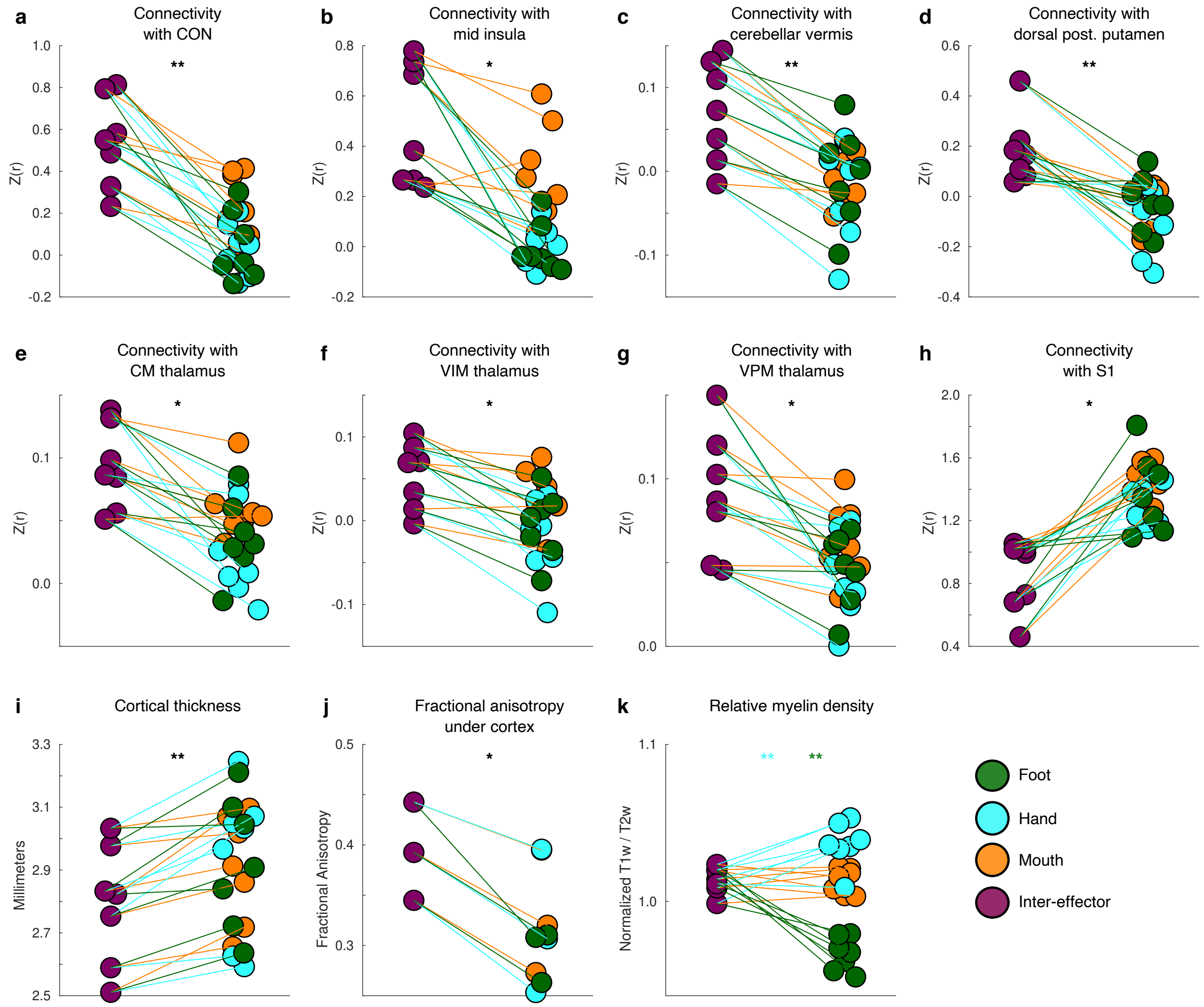


**Fig. S5| Functional connectivity and structural MRI metrics of motor cortex regions.** In each individual participant, measures derived from each of the foot, hand, mouth, and inter-effector motor regions. Colored lines connect the same participant's inter-effector and effector-specific regions for ease of comparison. **a,** Functional connectivity strength between M1 region and individual-specific cingulo-opercular network. **b,** Functional connectivity strength between M1 region and middle insula. **c,** Functional connectivity strength with Lobule VIIIa vermis of the cerebellum. **d,** Functional connectivity strength between M1 region and dorsal posterior putamen. **e-g,** Functional connectivity strength between M1 region and nuclei of the thalamus: **e,** Centromedian nucleus; **f,** Ventral Intermediate nucleus; **g,** Ventral Posteromedial nucleus. **h,** Functional connectivity strength between M1 region and adjacent postcentral gyrus (S1). **i,** Cortical thickness in M1 region. **j,** Fractional Anisotropy within 2 mm below cortex under M1 region. **k,** Intracortical myelin, indexed by the T1/T2 ratio and normalized across cortex, within cortex of M1 region. * $P < 0.05$; ** $P < 0.01$ ; *** $P < 0.001$.

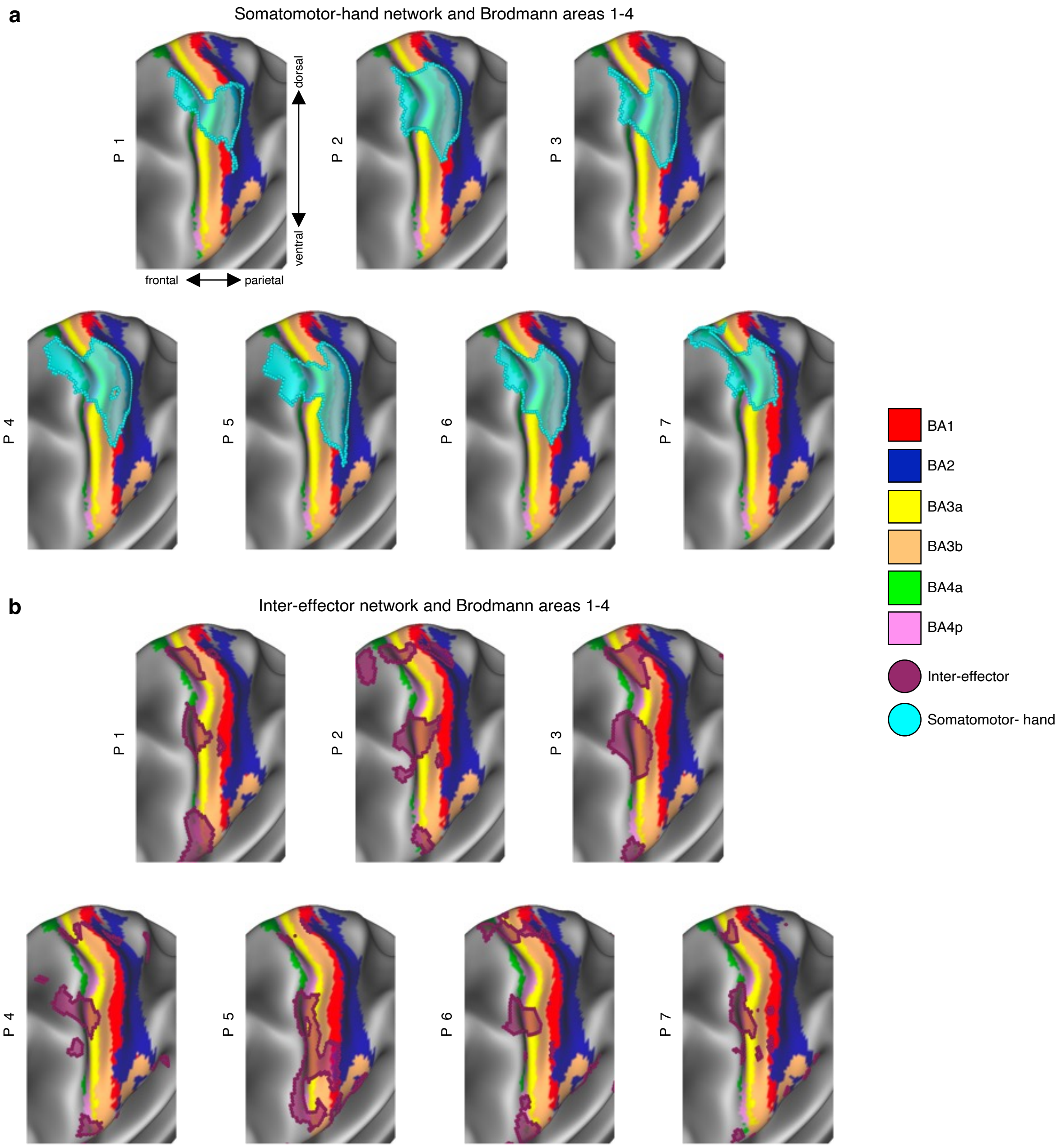


**Fig. S6| Inter-effector and effector-specific regions in pre- and postcentral gyrus.** In every participant, Brodmann Areas (BAs) in M1 (BAs 4a, 4p) and S1 (BAs 1, 2, 3a, 3b) are displayed on the cerebral cortex, tilted around the Y- and Z-axes to show S1. Overlaid are **a,** the somatomotor-hand region, and **b,** the inter-effector regions.

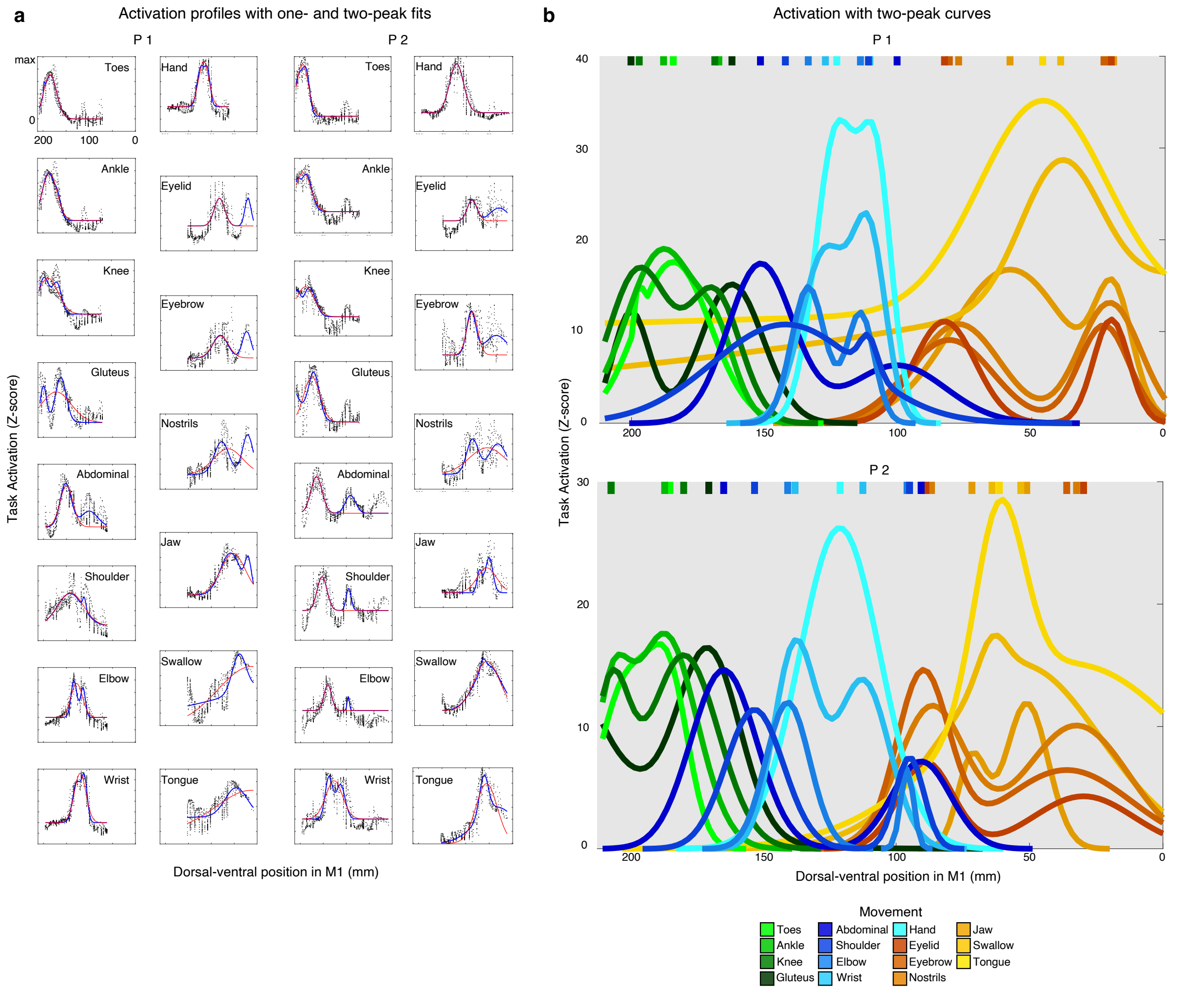


**Fig. S7| Concentric organization of task fMRI activations. a,** Movement-driven activation plotted against dorsal-ventral position within left hemisphere M1 for all movement tasks. For most movements, a two-peak curve (blue), modeled as a double-gaussian, fit the data better than a one-peak curve (red), modeled as a single-gaussian ($P < 0.001$). **b,** All modeled two-peak curves for P1 and P2. Most movements, except those by the distal-most body parts (toes, hand, tongue), showed a clear double peak. Peak locations (hashes on right) were arranged concentrically around the peaks of the distal-most body parts.

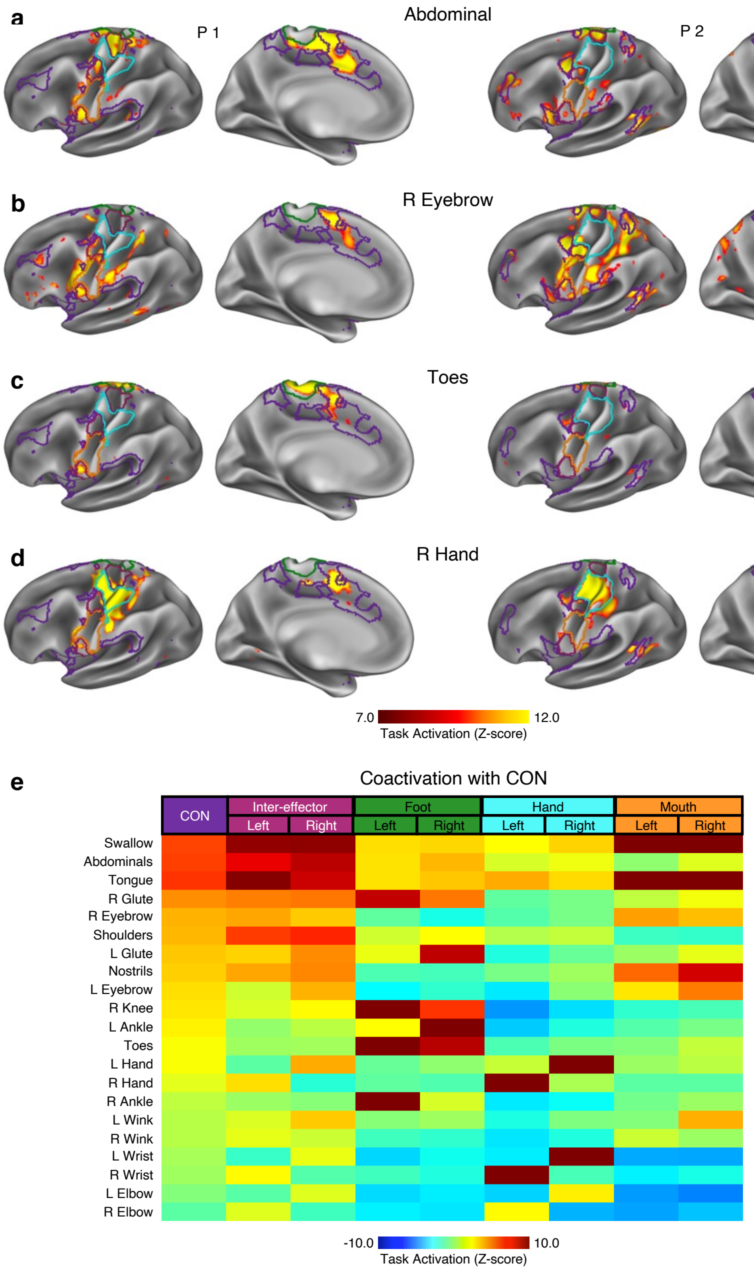


**Fig. S8| Effector-specificity of task fMRI activations.** In each participant, in the **a,** abdominal flexure task and the **b,** eyebrow raising task, the inter-effector regions and CON were active. By contrast, in **c,** toe and **d,** hand motion tasks, activation was much more specific to a single region of somatomotor cortex. **e,** Across tasks, the degree of CON activation was consistently similar to the activation of the inter-effector regions (correlation between CON and inter-effector activations: all $r > 0.81$, $P < 10^{-5}$), but not consistently to hand (CON vs hand: $r > 0.05$, $P < 0.82$) or foot (CON vs foot: $r > 0.33$, $P < 0.13$) regions, and more weakly to mouth regions (CON vs mouth: $r > 0.61$, $P < 0.003$). Illustrated activation values are averaged across participants and ordered based on CON activation.

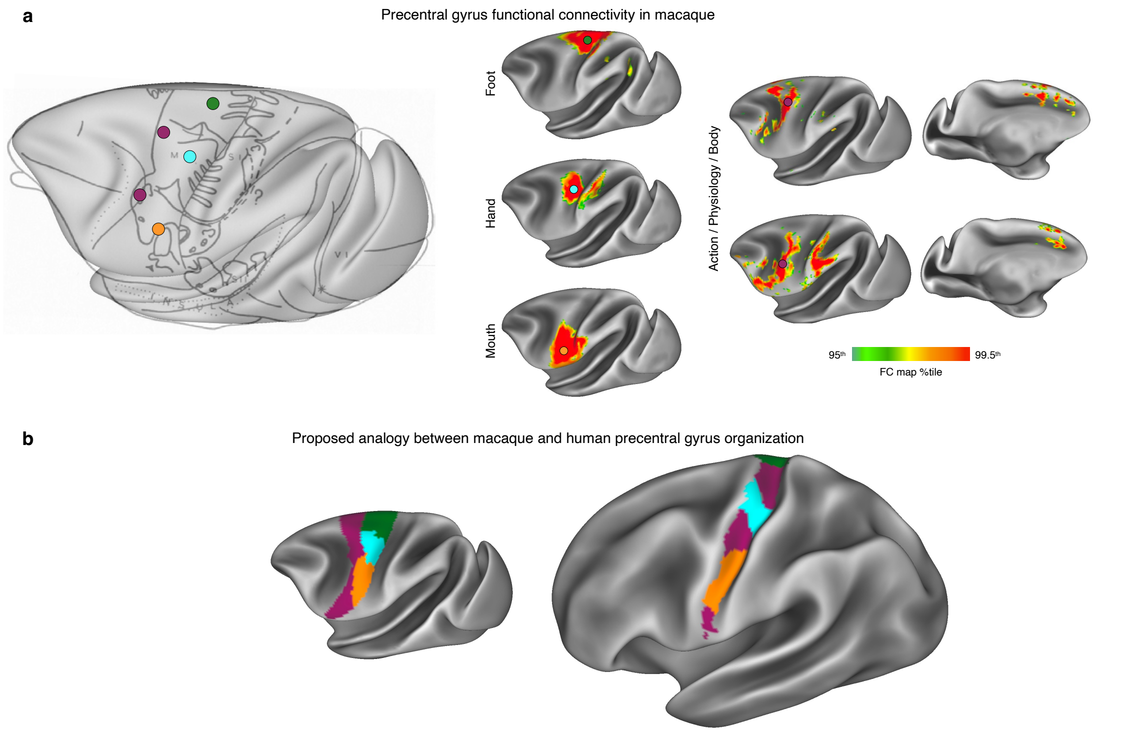


**Fig. S9| Functional connectivity in non-human primates. a,** Functional connectivity maps were seeded from points in precentral gyrus in a macaque. Left: seed locations are shown relative to the macaque "simiculus"[14]. Middle: maps seeded from posterior precentral gyrus locations corresponding to the foot (green), hand (blue), and mouth (orange) exhibited clear boundaries within M1 in the macaque, but no pattern of distributed connectivity interdigitated could be detected between the foot, hand, and mouth regions. Right: seeds in the anterior portion of precentral gyrus (maroon) exhibited strong connectivity distributed along a dorsal-ventral axis within anterior precentral gyrus, as well as with homologues of human CON regions, including inferior frontal gyrus, anterior inferior parietal cortex, and anterior dorsomedial prefrontal cortex. These anterior seeds corresponded to the body and neck, as well as to regions that are involved in complex actions[4] and which project to internal organs[10]. See Supplementary Movie 3 for complete mapping of motor connectivity in the macaque. This connectivity suggests that anterior motor regions in macaque may be analogous to the inter-effector regions in humans. **b,** Schematic illustrating proposed analogous regions in macaque (left) and in human (right). The foot region (green) is displaced in the human to the medial wall, while hand (cyan) and face (orange) regions maintained their relative positions. Anterior precentral regions in macaque (maroon) were displaced posteriorly in humans into the posterior bank of the precentral gyrus, interleaved between the foot, hand, and face regions.

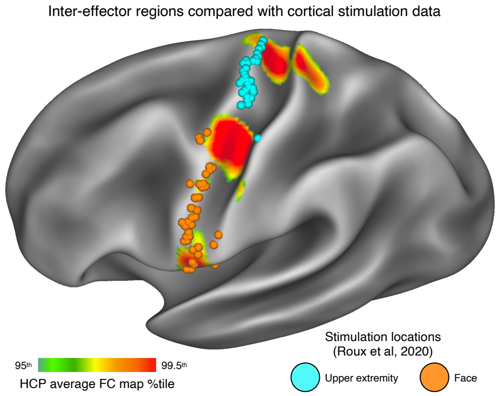


**Fig. S10| Inter-effector regions in human cortical stimulation data.** The map of inter-effector regions was compared with published movements evoked by direct cortical stimulation. Cortical map: functional connectivity is shown seeded from the central inter-effector M1 region and averaged across all subjects in the HCP dataset (see also Fig. S1c). Stimulation locations: MNI coordinates of stimulation location, and the resulting evoked movement, from 100 patients undergoing awake surgical brain mapping were reported in[53]. Each stimulation location evoking any movement was mapped to the nearest cortical vertex on a group average pial surface. Stimulation sites are colored according to whether they evoked facial movements (orange) or upper extremity movements (cyan). Stimulation sites evoking movement did not overlap with the central inter-effector region.

### Supplemental Discussion

#### Implications of a mind-body integration system for understanding and treating neurological disorders

The Mind-Body Interface includes two important thalamic nuclei that serve as deep brain stimulation (DBS) targets for clinical pathologies: the VIM for tremor (e.g., Essential Tremor[96], Parkinson's Disease[96,97]) and the CM for generalized seizures (e.g. Lennox-Gastaut Syndrome[98]). The mechanisms of action underlying these clinical effects are still debated, but it has been proposed that the anti-tremor effects of the VIM may be mediated by its connectivity to the cerebellum, and that the CM's role in arousal may explain its anti-epileptic effects.

Links with the MBI may be critical for the effects of neuromodulation on movement disorders. Tremor and arousal represent important aspects of integrative action control. Physiological tremor (~ 10 Hz) is thought to time and coordinate movements[67], while physiological arousal and CON engagement are observed as goal-directed activity begins[99].

Many types of tremors are intention- or goal-related. For example, essential tremor is absent or minimal at rest and is brought out by intentional movements. Further, most tremors disappear in sleep[100]. Finally, tremor and generalized seizures are global phenomena, not characterized by somatotopic specificity. Thus, the effects of VIM and CM DBS are consistent with the modulation of different aspects of whole-body action control, movement timing, and arousal.

Parkinson's disease (PD) may be most specifically related to dysfunction of MBI circuitry. PD symptoms cut across motor, physiological and volitional domains (e.g., postural instability, autonomic dysfunction, and reduced self-initiated activity, among many others[62]), mirroring MBI connections to regions relevant for postural control (cerebellar vermis), volition (dACC), and physiological regulation (insula) [6,7,57,58]. Work by Clinton Woolsey documented that direct stimulation in M1 of a PD patient temporarily eliminated his whole-body tremor and rigidity[49]. This effect is difficult to explain as a result of stimulation of effector-specific regions, but is fairly straightforward as a consequence of MBI stimulation. Neuronal death in the substantia nigra (SN) is one of the pathophysiological hallmarks of PD. Interestingly, the main target of SN projections is the dorsolateral putamen, which forms part of the Mind-Body Interface. Inter-effector regions are also strongly functionally connected to the cerebellar vermis, important for postural control and known to be structurally connected to M1 in NHP[101]. In addition, cortical projections important for coordinating physiology with action plans (i.e., blood pressure, orthostasis) primarily originate in CON and anterior M1[6,7]. Thus, if PD is indeed a network disease[102], a fitting candidate for the network most affected by the resulting degeneration is the Mind-Body Interface.
